# Paneth and Paneth-like cells undergoing necroptosis fuel intestinal epithelial cell proliferation following IFN-γ stimulation

**DOI:** 10.1101/2023.05.13.540666

**Authors:** Maria R. Encarnacion-Garcia, Raul De la Torre-Baez, Maria A. Hernandez-Cueto, Laura A. Velázquez-Villegas, Aurora Candelario-Martinez, Perla H. Horta-López, Armando Montoya-García, Gustavo Alberto Jaimes-Ortega, Luis Lopez-Bailon, Zayda Piedra-Quintero, Gabriela Carrasco-Torres, Marlon De Ita, Maria del Pilar Figueroa-Corona, José Esteban Muñoz-Medina, Magdalena Sánchez-Uribe, Marco Antonio Meraz-Ríos, Saúl Villa-Treviño, Francisco Garcia-Sierra, Bulmaro Cisneros, Michael Schnoor, Vianney F. Ortíz-Navarrete, Nicolás Villegas-Sepúlveda, Ricardo Valle-Rios, Oscar Medina-Contreras, Lilia G. Noriega, Porfirio Nava

## Abstract

The quality of life in patients with inflammatory bowel diseases (IBD) is strongly impaired. Alterations of intestinal epithelial homeostasis contribute to the development and establishment of IBD. Intestinal Paneth and Paneth-like cells produce and secrete luminal proteins sustaining epithelial homeostasis. Here we show that IFN-γ stimulates Paneth and Paneth-like cells degranulation that triggers the proliferation of intestinal epithelial cells (IEC) in a Wnt/*β*-catenin independent manner. Degranulation in Paneth and Paneth-like cells was mTORC1 and necroptosis dependent. Remarkably, lack of IFN-γ, inhibition of mTORC1, or impeding necroptosis reduces IEC proliferation cytokine-mediated. Our findings identify a new role for IFN-γ in stimulating IEC proliferation through inducing degranulation of Paneth and Paneth-like cells which is mTORC1 and necroptosis-dependent. In a mouse model of colitis, mTORC1 activation and necroptosis regulate Paneth and Paneth-like cell secretion. Furthermore, the colitogenic environment triggers PC metaplasia in the distal region of the large intestine to simulate cell proliferation.

**Highlights:** IFN-γ stimulates proliferation, *β*-catenin independent.

IFN-γ enhances mitochondrial activity and proliferation

IFN-γ regulates PC biogenesis.

mTORC1-dependent necroptosis mediates secretion in Paneth and Paneth-like cells.

## INTRODUCTION

Harmful luminal conditions constantly ravage IEC lining the gut. Old/damaged cells must be replaced by newly generated cells arising from stem cells residing in the crypts of Lieberkühn. Constitutive degranulation of Paneth (PC, Small intestine, Si) and Paneth-like cells (PLC, Large intestine, Li) stimulates epithelial cell proliferation (ECP), while luminal signals enhance epithelial cell death (ECD) to remove damaged/aged cells^1, 2^. Together, both processes maintain the number of IEC constant and the epithelial barrier functional.

Normal occurrence of ECD in the gastrointestinal tract of healthy individuals is mediated by apoptosis, anoikis, necroptosis, pyroptosis, and ferroptosis. Apoptosis and anoikis guarantee a clearance of damaged cells ensuring minimal risk of detrimental inflammation, and marginally disrupt epithelial barrier functions. In contrast, necrosis, necroptosis, and pyroptosis potentiate immune responses by releasing intracellular contents acting as damage-associated molecular patterns (DAMPs)^3–7^.

Interferon-gamma (IFN-γ) disrupts IEC homeostasis through altering several signaling pathways in the gut of IBD patients^8–10^. IFN-γ activates the serine/threonine kinase mammalian Target of Rapamycin (mTOR), a central core of two protein complexes, mTORC1, and mTORC2. mTORC1/C2 control various physiological processes such as cell growth/proliferation, development, and immunity^11^. In IEC, mTOR inhibits apoptosis, anoikis, and autophagy while promoting survival, differentiation, and proliferation^12–14^.

In addition to mTOR signaling, mitochondrial function can influence ECP, self-renewal, and ECD. Indeed, metabolic rewiring, characterized by increased respiration (high mitochondrial activity), occurs in stem cells undergoing proliferation and differentiation, while stem cell self-renewal encompasses low mitochondrial activity and increased glycolysis^15^. Additionally, overproduction of reactive oxygen species (ROS) due to mitochondrial hyperactivity promotes apoptosis in mature surface IEC^16^. Therefore, mTOR signaling impairment, mitochondrial dysfunction, and ROS overproduction can alter epithelial homeostasis and contribute to IBD development. However, regulation of these critical players by mucosal IFN-γ is partially understood.

Here we directly connected IFN-γ, mTOR, necroptosis, and ECP. Mechanistically, IFN-γ enhanced degranulation in PC and PLC to stimulate ECP in a *β*-catenin-independent fashion. Degranulation in PC and PLC entails mTORC1 activation and necrosome formation. Also, IFN-γ treatment resulted in mitochondrial hyperactivation and ROS overproduction. ROS, mTORC1, or necroptosis inhibition before IFN-γ stimulation prevented ECP and PC and PLC secretion. Additionally, we revealed that IFN-γ is constitutively secreted in Si. Furthermore, a colitogenic environment stimulated mitochondrial hyperactivation, ROS production, mTORC1 activation, necroptosis, and PC and PLC secretion. PC metaplasia also occurs during colitis. In conclusion, ECP following IFN-γ stimulation involves mTOR activation and necroptosis induction in PC and PLC.

## Material and methods

A detailed description and extended methodology and materials can be found as supplementary material.

## RESULTS

### Mitochondrial activity, ROS production, IEC proliferation, apoptosis, and necroptosis during the progression of DSS-induced colitis

Symptomatology evolution of Dextran Sulfate Sodium (DSS)-induced colitis reflects the gradual expression of factors involved in the pathogenesis, including mitochondrial and immune cell derived ROS^17, 18^. Therefore, we analyzed mitochondrial activity, ROS production, ECP, and ECD in mice exposed to DSS for 3 and 6 days to mimic different degrees of disease. DSS treatment progressively increased ECD (ACasp3 and the receptor-interacting serine-threonine kinase 3 (RIP3)) (**Figure 1A-B**) and reduced ECP (PCNA and pHist3) (**Figure 1C**), which resulted in colon length shortening (**Figure 1D and supplementary figure 1A**), extreme mucosal damage (**Figure 1E**), colonic crypt contraction (**Figure 1F**), and ulcer formation (**Figure 1G-H**). ROS concentrations increased following DSS treatment. However, ROS levels were higher at day 3 **(Figure 1I)** when epithelial damage and immune cell infiltration of ROS-producing PMN^18^ were less noticeable (**Figure 1A-G and supplementary figure 1B**). Therefore, we analyzed the cellular metabolism as a possible source of ROS through investigating oxygen consumption rate (OCR) at different states of mitochondrial respiration in isolated mitochondria harvested from ileum and colon of DSS-treated mice. As shown in **figure 1J**, basal mitochondrial OCR significantly increased after 3d of DSS treatment in the intestinal mucosa. However, after 6d of treatment, mitochondrial respiration was like control animals. Likewise, basal, maximal, and ATP-production associated respiration, and proton leak increased in Li mitochondria harvested from mice treated with DSS for 3d and was akin to control animals after 6d of treatment. Similar results were obtained in Si (**Supplementary figure 1C-E**). Thus, high mitocondrial activation and ROS overproduction simultaneously with increased ECP, modest ECD, marginal PMN infiltration and negligible epithelial damage occurred after 3d of DSS treatment. In contrast, exacerbated PMN infiltration, reduced ECP, augmented ECD and major mucosal erosion associated with intermediate levels of ROS and normal mitochondrial activity at 6d of DSS-treatment.

**Figure 1.**
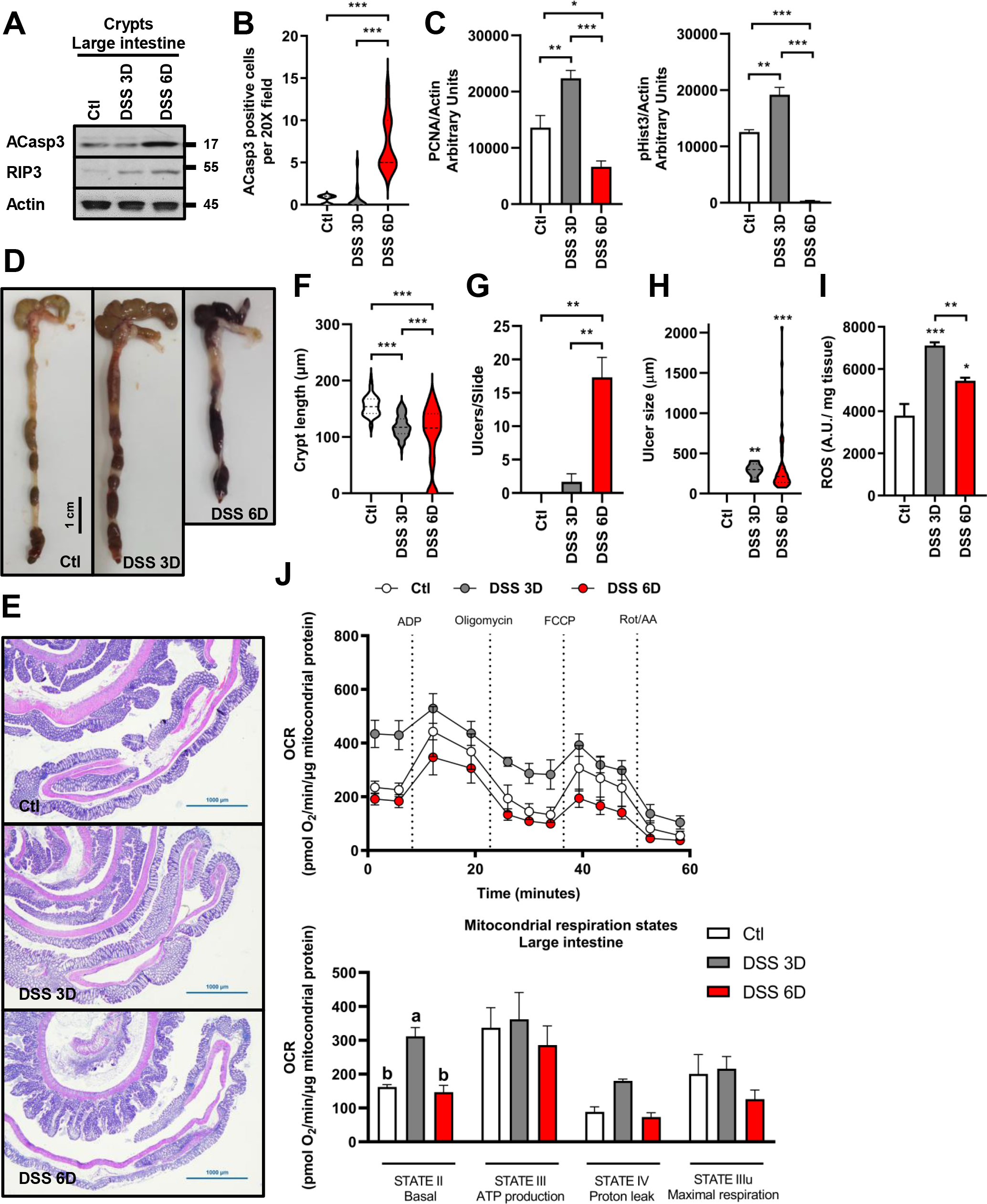
**Mitochondrial activity, apoptosis, and necroptosis in the colitic gut.** A) ACasp3 and RIP3 were evaluated by western blotting in isolated crypts of Li harvested from control and 2.5% DSS-treated animals. DSS treatment was carried out for 3 and 6d. Actin was used as loading control. n=6 independent experiments. B) Graph representing the quantification of ACasp3^+^ cells per high magnification field (20X) in the colonic mucosa of control and 2.5% DSS-treated mice. DSS treatment was carried out for 3 and 6d. 6 different animals per condition were evaluated. C) PCNA and pHist3 were evaluated by western blotting in isolated crypts of large intestine harvested from control and 2.5% DSS-treated mice. DSS treatment was carried out for 3 and 6d. Values were normalized to actin. Densitometric analysis of 4 independent experiments is shown. D) Representative pictures of the colon and H&E staining (E) from the colonic mucosa from control and 2.5% DSS-treated mice. Colon tissue was harvested after 3 and 6d of treatment. Bar= 1000µm. n=6 animals per group. F) Crypt length was analyzed in the colonic tissue of control and 2.5% DSS-treated mice. Colon tissue was harvested after 3 and 6d of treatment. Graph represents a pool of 100 crypts analyzed in 6 animals. Mucosal damage indicated by the number (G) and size of the ulcers (H) present in H&E- stained histological sections of colonic tissue of control and 2.5% DSS-treated mice is graph. 3 slides per animal were used for evaluation. n=6. I) Histogram representing ROS quantification in large intestine obtained from control and 2.5% DSS-treated animals. CellROX™ assay was performed in colonic tissue of mice treated with 2.5% DSS for 3 and 6d. n=3. J) Oxygen consumption rate (OCR) in isolated mitochondria from colonic tissue of control and DSS (2.5%)- treated mice after 3 and 6d of treatment. OCR was measured at basal conditions and following ADP, oligomycin, FCCP and antimycin A/rotenone injection. Respiration states were calculated by subtracting OCR values after antimycin A/rotenone administration. n=4-8 mice per group. Data are shown as mean ± SEM and are pooled from 3 independent experiments. P values were calculated using one-way analysis of variance with the Tukey post hoc test (B, C, F, G, H, I). *p< 0.05; **p < 0.01; ***p < 0.001.

### ROS scavenging attenuates DSS-induced colitis

DSS-treatment increased ROS, RIP3, and ACasp3 in colonic IEC, while diminishing EdU, pHist3 and PCNA **(Figure 2A-C)**. Likewise, *in vitro* DSS administration resulted in ROS overproduction, augmented ECD and reduced ECP **(Supplementary figures 2A-B).** To evaluate which of these effects can be directly attributed to ROS, SW480 cells were treated with hydrogen peroxide (H2O2)^19^. Remarkably, H2O2 reduced ECP and stimulated ECD (**Supplementary figures 2C**). Therefore, we evaluated ECP and ECD in the colonic mucosa of DSS-treated mice simultaneously administered with the ROS scavenger, N-acetylcysteine (NAC)^20^. NAC administration further enhanced body weight loss, Disease Activity Index (DAI), and immune cell infiltration in the colitic mucosa of DSS-treated mice (**Supplementary figures 2D-F**). However, prevented colon length shortening (**Figure 2D**), reduced the histological score (**Figure 2E**), diminished ECD and restored ECP (**Figure 2F**) after 6d of DSS treatment. However, in animals treated with DSS for 3d the ROS scavenger attenuated ECP and apoptosis without affecting necroptosis (**Supplementary figures 2G**). These results suggest that the source and amount of mucosal ROS differentially influence ECP and ECD.

**Figure 2.**
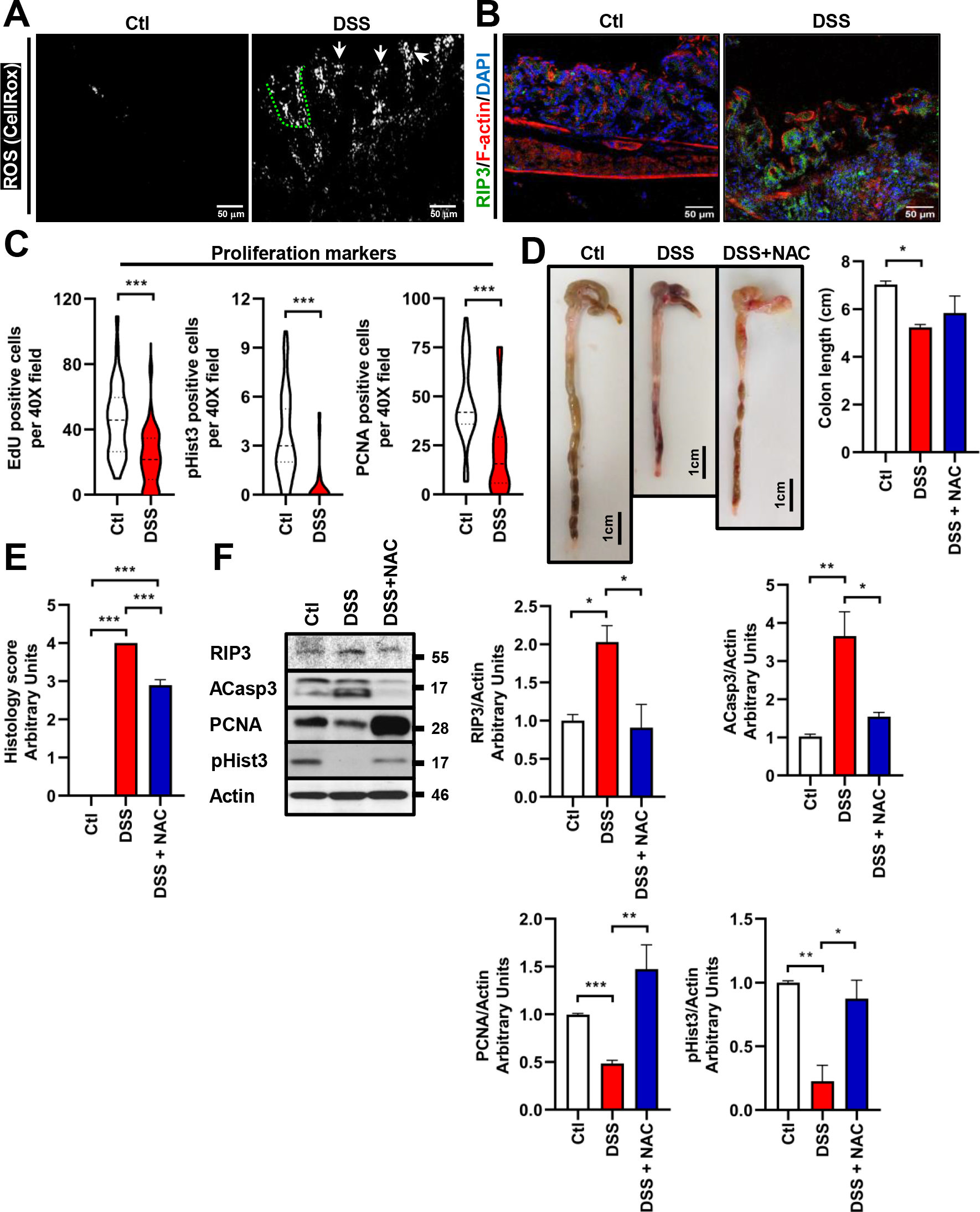
ROS scavenging attenuates DSS-induced damage in the colitic mucosa. A) ROS was evaluated in the colonic mucosa of control and colitic mice. C57BL/6J mice received water alone or 2.5% DSS dissolved in drinking water for 6d. CellROX™ (white) was used to detect ROS in 10< m cryosections of colonic tissue. Arrows=surface IEC. Discontinuous green line limits colonic crypt. Bar=50*μ*m. A representative image of n=6 independent experiments is shown. B) Receptor-interacting serine/threonine-protein kinase (RIP)3 (green) and F-actin (red) were evaluated by immunofluorescence staining in colonic cryosections of control and colitic mice. 2.5% DSS dissolved in drinking water was administered to Wt mice for 6d. Nuclei=blue. Bar=50*μ*m. A representative image obtained from 6 different animals is displayed. C) IEC proliferation markers EdU, pHist3, or PCNA were evaluated in colonic cryosections of control and colitic mice. n=6 different animals per condition were evaluated. Graphs displaying the number of proliferating IEC per high magnification field (40X) are shown. D) Representative pictures of the large intestine from water/PBS, 2.5% DSS/PBS, 2.5% DSS/N-acetylcysteine (NAC)-treated mice are shown. Graph displaying colon length is shown. Colon tissue was harvested from euthanized mice after 6d of treatment. Mice received a daily i.p. injection of vehicle alone (PBS) or NAC (100 mg/Kg). Bar= 1cm. n=6 animals per group. E) Histology scoring of the colonic tissue harvested from mice treated with water/PBS, 2.5% DSS/PBS, 2.5% DSS/NAC-treated mice for 6d. Histology score is expressed in arbitrary units. Colon tissue was harvested from euthanized mice after 6d of treatment. Mice received a daily i.p. injection of vehicle alone (PBS) or NAC (100 mg/Kg). n=6 animals per group. F) RIP3, ACasp3, PCNA and pHist3 were evaluated by western blotting whole colonic mucosa obtained from water/PBS, 2.5% DSS/PBS, 2.5% DSS/NAC-treated mice. Actin was used as loading control. Graphs showing mean densitometric values in the biological samples are presented. Representative blot of n=3 independent experiments (2 animals/group) is shown. Data are shown as mean ± SEM and are pooled from 3 independent experiments. P values were calculated using *t*-test (C), one-way analysis of variance with the Tukey post hoc test (D-F). *p< 0.05; **p < 0.01; ***p < 0.001.

### IFN-γ enhances mitochondrial activity and IEC proliferation in the gut

ECD ROS- mediated has been thoroughly studied during colitis^21, 22^ Nonetheless, the role of ROS in the initiation of ECP remains unknown. Monitoring changes in the intestinal mucosa of DSS-treated mice is complicated as the disease develops^23^ but, intraperitoneal administration of IFN-γ, is a reliable and controllable method that partially recapitulates biochemical symptoms of colitis^24^. Thus, we investigated a putative link between ROS formation, mitochondrial activation and ECP in the gut of IFN-γ-treated mice. IFN-γ was administered at 4 and 18h to analyze early and late effects of the cytokine. As shown in **figure 3A,** mitochondrial basal and ATP-production OCR increased at 4h post-IFN-γ stimulation, and the effect was attenuated after 18h of treatment. Mucosal ROS and ECD (ACasp3 and RIP3) gradually and sustainably increased in IEC following IFN-γ stimulation (**Figure 3B-D**). Additionally, long-time exposure to IFN-γ enhanced PMN infiltration (**Data not shown**), reduced ECP (**Figure 3D-E and supplementary figure 3A**) and shortened the colonic crypts and collapsed its lumen (**Figure 3F-G).** Unexpectedly, 4h of IFN-γ treatment stimulated ECP (**Figure 3D-E and Supplementary figure 3B**), enlarged the colonic crypts without inducing cytoarchitectural modifications (**Figure 3F- G**) or the recruitment of PMN into the tissue (**Data not shown**). Similar effects were observed in Si, except that the cytokine unaffected the crypt-villus axis length (**Supplementary figure 3C-H**). Thus, short-term stimulation with IFN-γ enhances mitochondrial activation, ROS formation and ECP in the gut.

**Figure 3.**
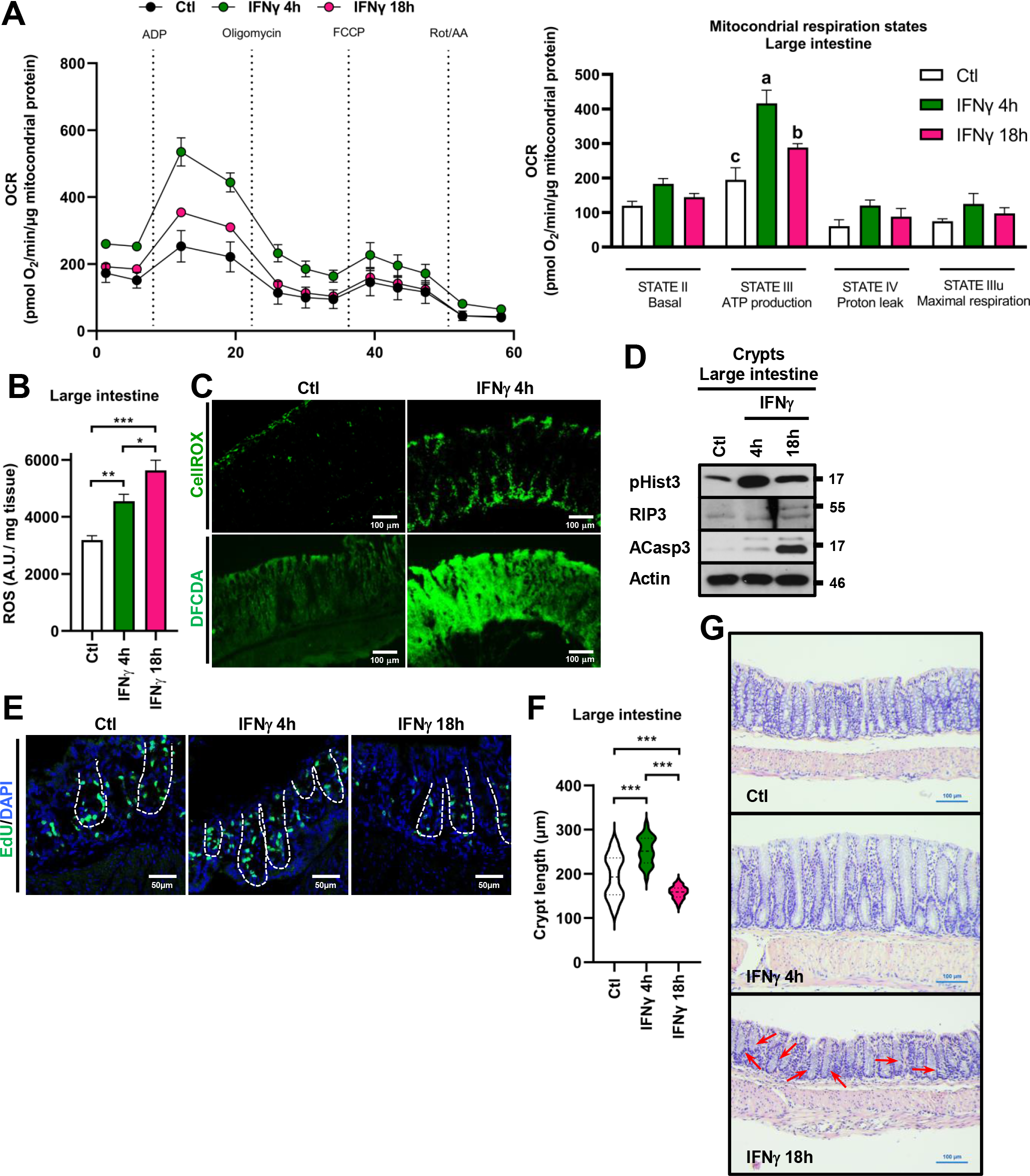
IFN-γ transitorily increases mitochondrial activity and cell proliferation of IEC. A) Oxygen consumption rate (OCR) in isolated mitochondria from colonic tissue of C57BL/6J mice injected i.p. with vehicle (Mouse Serum Albumin, MSA) or IFNγ (2.5 mg/kg) for 4 or 18h post-injection. OCR was measured at basal conditions and following ADP, oligomycin, FCCP and antimycin A/rotenone injection. Respirations states were calculated by subtracting OCR values after antimycin A/rotenone administration. n=4-8 mice per group. B) ROS quantification in large intestine of C57BL/6J mice injected i.p. with vehicle (Mouse Serum Albumin, MSA) or IFN-γ (2.5 mg/kg). Colonic mucosal tissue was collected 4h or 18h post- cytokine injection and incubated for 30 minutes with CellROX™ reagent. Fluorescence was analyzed in a Fluoroskan FL Microplate Fluorometer. n=6. C) ROS distribution was evaluated in the colonic mucosa of C57BL/6J mice injected i.p. with MSA or IFN-γ (2.5 mg/kg) using fluorescence microscopy. Mice were euthanized 4h post-cytokine injection. CellROX™ (green) or 2’,7’– dichlorofluorescein diacetate (DFCDA, green) were used for ROS detection. Bar=100*μ*m. n=6. A representative image is provided. D) pHist3, RIP3 and ACasp3 were evaluated by western blotting colonic isolated crypts obtained from C57BL/6J mice injected i.p. with MSA or IFN-γ (2.5 mg/kg). Mice were euthanized 4 or 18h post-cytokine injection. Actin was used as loading control. n=6 independent experiments. E) EdU (green) incorporation was analyzed by confocal microscopy in colonic cryosections of C57BL/6J mice i.p. injected with MSA or IFN-γ (2.5 mg/kg). Mice were euthanized 4 or 18h post-cytokine injection. Representative images are shown. Nuclei=blue. Discontinuous white line limits the colonic crypt. Bar=50*μ*m. F) Crypt length was analyzed in the colonic tissue of mice i.p. injected with MSA or IFN-γ (2.5 mg/kg). Mice were euthanized 4 or 18h post-cytokine injection. Graph represents a pool of 100 crypts analyzed in 6 animals. G) Representative H&E staining from colonic mucosa of C57BL/6J mice injected i.p. with MSA or IFN-γ (2.5 mg/kg). Mice were euthanized 4h or 18h post injection. Arrow indicates collapsed luminal crypt. Bar= 100µm. Representative images are displayed. n=6 animals per group. Data are shown as mean ± SEM and are pooled from 3 independent experiments. P values were calculated using one-way analysis of variance with the Tukey post hoc test (B, F). *p< 0.05; **p < 0.01; ***p < 0.001.

### IFN-γ-mediated cell proliferation is mTOR dependent

Long-term effects IFN-γ- mediated are broadly documented^25–27^. Nevertheless, the machinery spurring ECP downstream of the cytokine remains poorly understood. Wnt/*β*-catenin signaling drives ECP in the gut. However, IFN-γ stimulation rapidly inhibited *β*-catenin signaling as observed by the reduction of phospho-LDL receptor related protein-6 (LRP6) and EGFP in the gut of the Lgr5-EGPF reporter mice (**Figure 4A and supplementary figure 4A**). IFN-γ treatment minimally affected the amount and localization of *β*-catenin or its active form (ABC) (**Supplementary figure 4A-C**). Surprisingly, 4h of IFN-γ treatment augmented the stem cell marker Sca-1 in the intestinal mucosa (**Figure 4B**). To verify those findings, we analyzed the expression of *Lgr5* and *Olfm4*, another intestinal epithelial stem cell marker, in isolated crypts from Si and Li of Wt mice exposed to IFN-γ. As shown in **figure 4C-D**, IFN-γ administration for 4h reduced *Lgr5* expression but induced *Olfm4* and the expression of both markers was reduced after 18h of cytokine treatment. Other biological events related to ECP were also enhanced after 4h of IFN-γ-treatment, such as the reduction of intraepithelial cell adhesion uncovered by the decrease of EpCAM (epithelial cell adhesion molecule) (**Supplementary figure 4D-E**)^28^. Thus, these data suggest that IFN-γ rapidly promotes a proliferative phase that could foster the generation of newborn stem cells.

**Figure 4.**
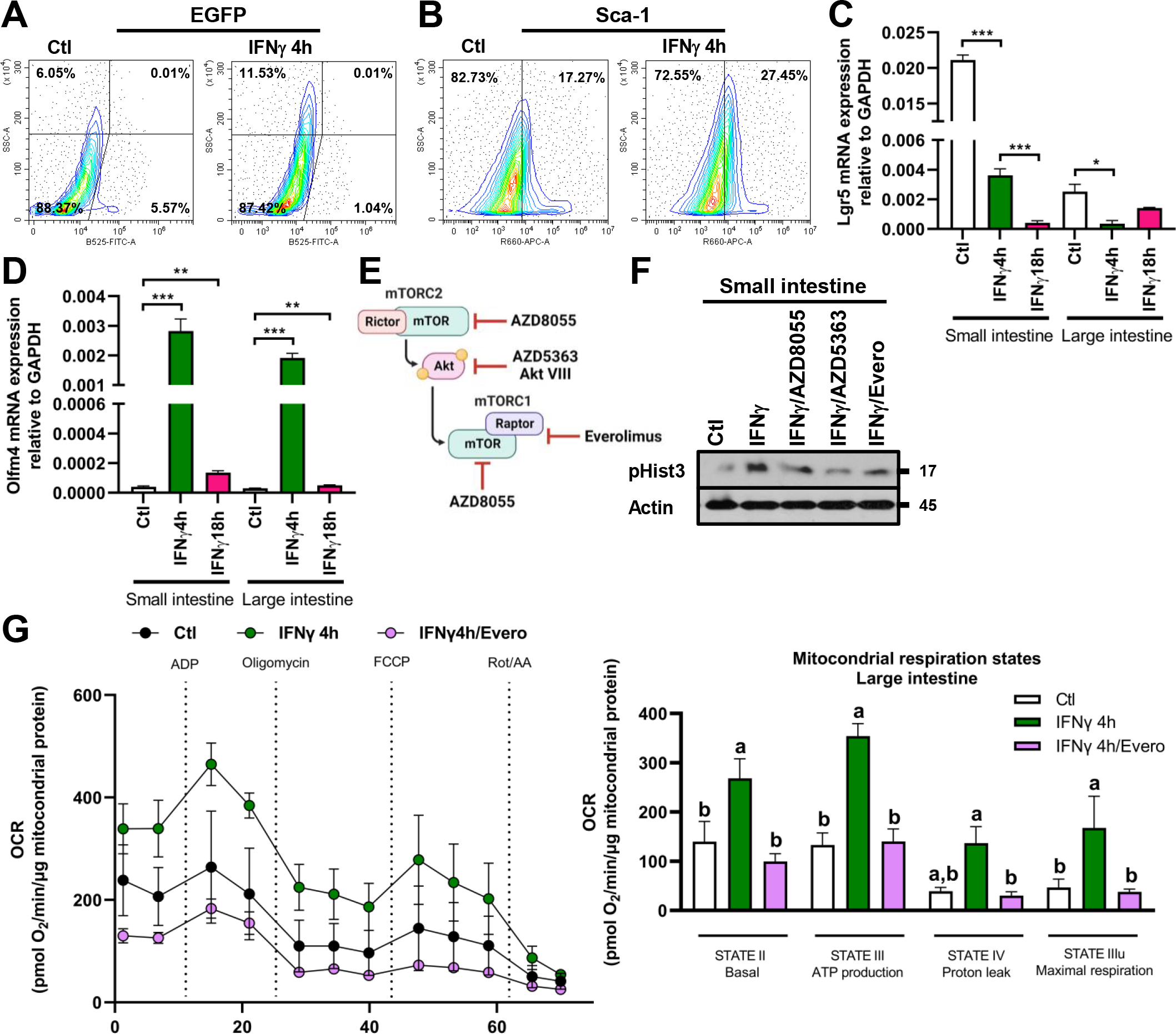
IFN-γ increases cell proliferation mTOR-mediated while inhibiting β- catenin transcriptional activity. Flow cytometric identification of EGFP+ (A) and Sca-1+ (B) cells in preparations pooled from ileum of Lgr5-EGFP-IRES-creERT2 “knock-in” reporter mice. Cells were labeled with Sca-1-FITC. Values represent the mean percentage of events falling within their respective quadrants. Small intestine was harvested from mice treated with MSA i.p. (Ctl) or IFN-γ (2.5 mg/kg, i.p.) for 4h. n=4. *Lgr5* (C) and *Olfm4* (D) mRNA expressions were analyzed in small and large intestine by qPCR. Crypt isolation was carried out from mice treated with MSA i.p. (Ctl) or IFN-γ (2.5 mg/kg, i.p.) for 4 or 18h. *Lgr5* and *Olfm4* expression was normalized to GAPDH. n=3. E) mTOR (mammalian target of rapamycin) protein is the central core of mTORC1 and mTORC2. AZD8055 is an ATP-competitive mTOR inhibitor. AZD5363 and AKT Inhibitor VIII potently inhibit all Akt isoforms (Akt1/Akt2/Akt3). The rapalog, Everolimus, suppress mTORC1 activity. F) pHist3 was evaluated by western blotting and IHC in IEC of the small intestine of C57BL/6J mice injected i.p. with MSA/DMSO (Ctl), IFN-γ (2.5 mg/kg)/DMSO, IFN-γ/AZD8055 (25 mg/Kg, i.p.), IFN-γ/AZD5363 (30 mg/Kg, i.p.) or IFN-γ/Everolimus (1 mg/Kg, i.p.). Mice were euthanized 4h post-cytokine injection. Inhibitors or carrier were administered 30 min before cytokine injection. Actin was used as loading control. n=4 independent experiments. G) Oxygen consumption rate (OCR) in isolated mitochondria from colonic tissue of C57BL/6J mice injected i.p. with MSA (Ctl), IFNγ (2.5 mg/kg), or IFNγ/Everolimus (1 mg/Kg, i.p.) after 4h of treatment. OCR was measured at basal conditions and following ADP, oligomycin, FCCP and antimycin A/rotenone injection. Respiration states were calculated by subtracting OCR values after antimycin A/rotenone administration. n=4-8 mice per group.

Maintenance of intestinal stem cell progenitors Olfm4^+^ is mTOR-mediated and Wnt- independent^14^. Hence, we investigated if mTOR signaling was driving ECP following IFN-

γ treatment. Then, Wt mice were administered with mTOR inhibitor (AZD8055), Akt inhibitor (AZD5363, Akt inhibitor VII) or mTORC1 inhibitor (Everolimus) 30 min prior IFN-γ administration (**Figure 4E**). Inhibition of mTOR, Akt or mTORC1 averted the increase in ECP induced by the cytokine in Si and Li (**Figure 4F and Supplementary figure 4F**). Furthermore, Everolimus efficiently inhibited the increment on mitocondrial respiration triggered by IFN-γ (4h) in Si and Li (**Figure 4G and Supplementary figure 4G**), demonstrating a role of mTORC1 in the process.

### IFN-γ requires mTORC1 activation and necroptosis to stimulate cell proliferation through PC

PC (**Figure 5A**) degranulation mTORC1-mediated^29^ instigates ECP^30^ and IFN-*γ* stimulates PC secretion^31^. Then, mTORC1-activation in PC could facilitate ECP IFN-γ-mediated. Consistent with our theory, in control conditions, phospho-ribosomal protein S6 (pS6) staining revealed mTORC1 activation in few cells at the bottom of the crypts of Lieberkühn, but IFN-γ rapidly activated mTORC1 in IEC at the crypt bottom, including in PC. However, by 18h of cytokine spurring active mTORC1 was detected mainly in surface IEC (**Figure 5B-C**).

**Figure 5.**
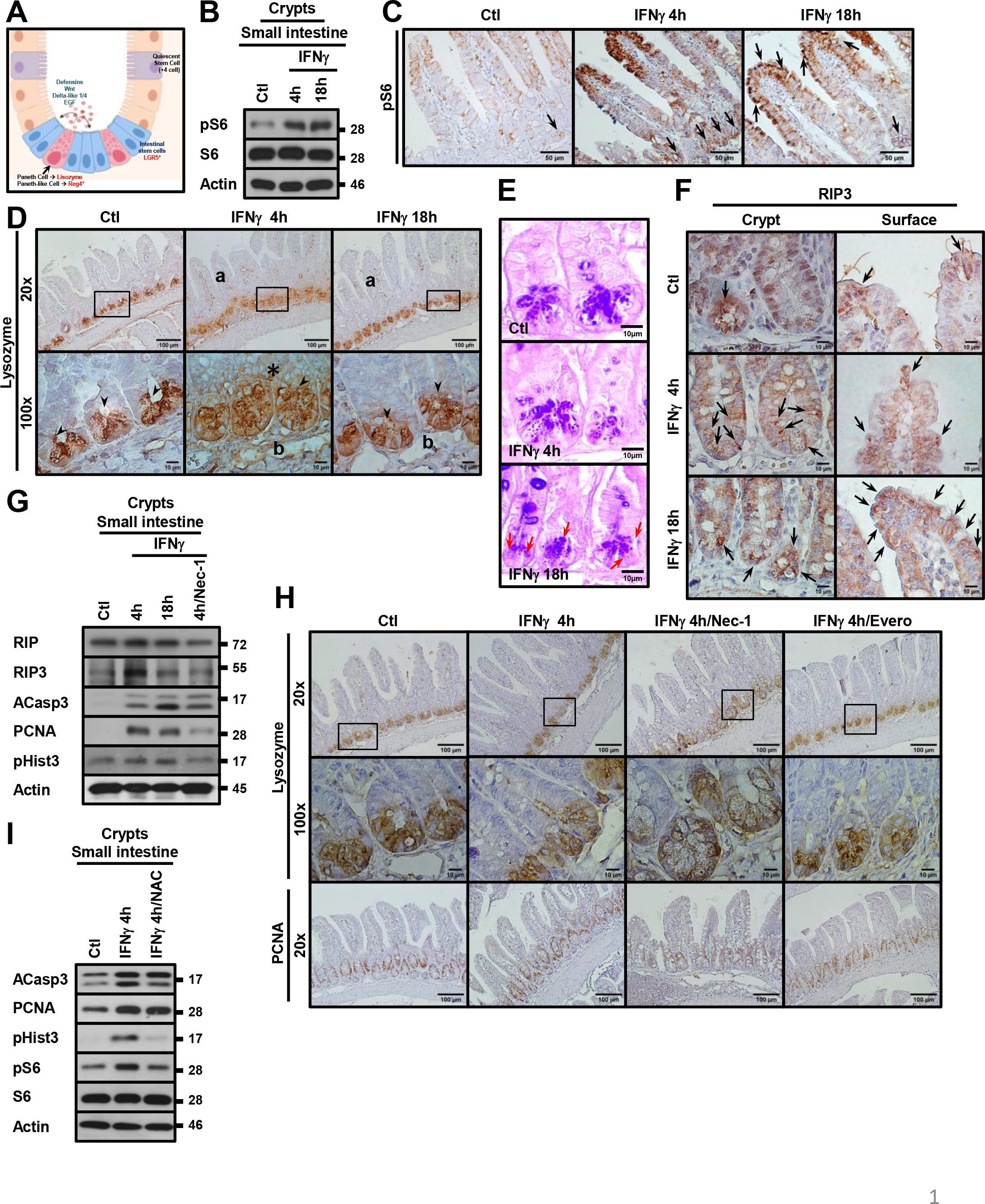
Necroptosis mTORC1-mediated mediates PC degranulation following IFN-γ stimulation. A) Paneth and PLC at the bottom of the intestinal crypts maintain the progenitor niche by secreting defensins and growth factors that target neighbored cells. pS6 and S6 were assessed in IEC of the small intestine of C57BL/6J mice injected with MSA (Ctl) or IFN-γ (2.5 mg/kg) by western blotting (B) and IHC (C). Mice were euthanized 4 or 18h post-cytokine injection. Bar=50 μm. Actin was used as loading control. n=4. D) Lyzozyme-1 distribution was analyzed by IHC in 4 μm paraffin embedded sections of small intestine from control and IFN-γ-treated mice. Mice were euthanized 4 or 18h post-cytokine injection. Lower panel= amplification of square region marked in upper panel. Arrowheads= crypt-lumen lyzozyme-1. a=non-epithelial cells Lyzozyme-1+. Asterisk= non-PC lyzozyme-1+. b= PC lyzozyme-1+. n= 6 animals per group. Bar=100*μ*m and 10*μ*m. E) Micrograph of PC of control and IFN-γ-treated mice. H&E staining was carried out in 4 μm paraffin embedded sections of small intestine. Mice were euthanized 4 or 18h post- cytokine injection. Arrows= atrophic PC. Bar=10 μm. F) RIP3 was evaluated by IHC in 4 μm paraffin embedded sections of small intestine from control and IFN-γ-treated mice. Mice were euthanized 4 or 18h post-cytokine injection. Upper panel= Amplification of the villus surface. Lower panel= Amplification of the crypts of Lieberkühn. Arrows= necrosomes RIP3+. n= 6 animals per group. Bar=10 μm. G) RIP, RIP3, ACasp3, PCNA and pHist3 were evaluated by western blotting isolated crypts of small intestine obtained from C57BL/6J mice injected i.p. with MSA (Ctl), IFN-γ (2.5 mg/kg) or IFN-γ/Necrostatin-1 (Nec-1, 4.5 mg/Kg, i.p.). Necrostatin-1 was administered i.p. 30 min before cytokine stimulation. Mice were euthanized 4h or 18h post-cytokine injection. Actin was used as loading control. H) Lyzozyme-1 and PCNA were evaluated by IHC in 4 μm paraffin embedded sections of small intestine from control, IFN-γ, IFN-γ/Everolimus and IFN-γ/Necrostatin-1-treated mice. Mice were euthanized 4h post-cytokine injection. Necrostatin-1 and Everolimus were administered i.p. 30 min before cytokine stimulation. 20x = low magnification. 100x= High magnification. Black square= Magnified areas. n= 6 animals per group. Bar= 100*μ*m and10*μ*m. I) ACasp3, PCNA, pHist3, pS6 and S6 were evaluated by western blotting isolated crypts of small intestine obtained from C57BL/6J mice injected i.p. with MSA (Ctl), IFN-γ (2.5 mg/kg) or IFN-γ/N-acetylcysteine (NAC). NAC (100 mg/Kg) was administered i.p. 30 min before cytokine stimulation. Mice were euthanized 4 or 18h post-cytokine injection. Actin was used as loading control.

Considering these results, we traced PC function through lysozyme-1^32^ (**Supplementary figure 5A**). IFN-γ treatment transiently induced lysozyme-1 accumulation in enterocytes (**Supplementary figure 5B**) despite of apparently inhibiting its expression (**Supplementary figure 5C**). Furthermore, in control conditions lizozyme-1 was present in cytosolic granules of PC with small amounts of the protein invading the crypt-lumen (**Figure 5D, arrowhead**). However, at 4h post-cytokine stimulation, lysozyme-1 appeared less granular and more diffuse in the cytosol with a large amount of the molecule found in the crypt-lumen (**Figure 5D, a**). Additionally, the molecule was detected in the lateral membrane of neighboring cells and in the cytosol of IEC lysozyme-1- (**Figure 5D, Asterisk**). By 18h, lysozyme-1+ granules were formed again, and the molecule in the crypt-lumen reduced (**a**). Importantly, submucosal cells lysozyme-1+ were frequently discovered after cytokine administration implicating synthesis of the protein in immune cells (**Figure 5D, b**). Histological analysis revealed healthy-looking PC in control, and after 4h of IFN-γ treatment (**Figure 5E**). However, shrink/damaged PC were detected following 18h of cytokine stimulation (**Figure 5E, red arrows**).

mTORC1 stimulates necroptosis in IEC, to consent the release of intracellular contents^33, 34^ and could therefore be involved in PC degranulation. Consequently, we evaluated the clustering of RIP3**^+^** in IFN-γ-treated mice. In control conditions, IEC along the crypt-axis displayed cytosolic RIP3 and few crypt-base and villus-surface cells exhibited amyloid microfilaments RIP3^+^ (**Figure 5F, arrows**). After 4h of IFN-γ administration, the number of IEC necrosomes+ augmented at the crypt base, while surface IEC necrosomes+ remained unchanged. By 18h of IFN-γ stimulation, IEC necrosome^+^ diminished at the crypt base and surface IEC necrosome+ augmented (**Figure 5F, arrows**). Consequently, we evaluated the effect of necrostatin-1, a RIP1/RIP3 inhibitor^35^, in the induction of ECP IFN-γ-mediated. By western blotting, RIP and RIP3 augmented at 4h but not by 18h of IFN-γ treatment and necrostatin-1 inhibited the increment induced by the cytokine (**Figure 5G**). Necrostatin-1 minimally affected Casp-3 activation demonstrating the specificity of the treatment. Furthermore, necrostatin-1 prevented the release of lysozyme-1 into the lumen of the crypts Lieberkühn and impeded the up-rise of the proliferation markers PCNA y pHist3 after 4h of IFN-γ-treatment (**Figure 5G-H**). Unsurprisingly, Everolimus administration yielded similar results to necrostatin-1 (**Figure 4F and 5H**) and prevented the crypt-lumen collapse and the PC morphological changes induced by 18h of IFN-γ treatment (**Supplementary figure 5D**).

In IEC, low concentrations of ROS upregulated ECP, necroptosis and mTORC1 signaling (**Supplementary figure 5E**). Furthermore, IFN-γ stimulated mitochondrial activity mTORC1 and ROS-dependent but necroptosis independent (**Supplementary figure 5F**). Therefore, we evaluated ECP, ECD, and mTORC1 activation in the colonic mucosa of mice treated with IFN-γ or IFN-γ/NAC. As expected, IFN-γ administration stimulated ECP, apoptosis, and mTORC1 activity in isolated crypts of Wt mice and scavenging ROS through NAC prevented those processes (**Figure 5I)**, confirming that ROS stimulate ECP, ECD, and mTORC1 signaling downstream of IFN-γ. NAC administration cannot prevent RIP3 upregulation induced by the cytokine (**Data not shown**).

PLCs expressing regenerating islet-derived family member 4 (Reg4^+^) control IEC homeostasis in the colon (**Supplementary figure 5A).** Thus, we evaluated mTORC1 activation and necroptosis in the base of the colonic crypts where PLCs reside^36^. Like the Si, 4h of IFN-γ increased pS6 and the number of assembled necrosomes in colonocytes along the whole crypt-surface axis including IEC at the crypt base (**Supplementary figure 5G-I**). However, by 18h, these changes were mainly observed at the crypt surface (**Supplementary figure 5G**). Furthermore, IFN-γ enriched Reg4 in colonocytes and the effect lasted for over 24h period (**Supplementary figure 5J**). Additionally, intraperitoneal administration of IFN-γ stimulated the secretion of Reg4 into the intestinal lumen (**Data not shown**). The results suggest that IFN-γ induces degranulation of PC and PLC by inducing necroptosis mTORC1-dependent.

### Constitutive secretion of IFN-γ in the small intestine modulates PC function

Indisputably, IFN-γ affects IEC homeostasis and PC and PLC function. Intraepithelial and lamina propria T cells spontaneously release IFN-γ in the gut a function impaired in mice lacking class I-restricted T cell-associated molecule (CRTAM)^37–39^. Thus, we evaluated IFN-γ secretion in Wt and CRTAM-deficient mice. In the beginning, we discovered that Wt mice yielded ∼100 times more IFN-γ in the Si than in Li, with a constitutive secretion of the cytokine occurring only in Si (**Figure 6A)**. IFN-γ was undetectable in the Si of CRTAM^KO^ animals. Surprisingly, IFN-γ levels were comparable amongst the colon of Wt and CRTAM^KO^ mice and the cytokine was released in the Li of CRTAM^KO^ mice (**Figure 6A**). Because of these results, we evaluated mTORC1 activation, apoptosis, necrosome formation and lysozyme-1 in the Si of Wt and CRTAM^KO^ mice.

**Figure 6.**
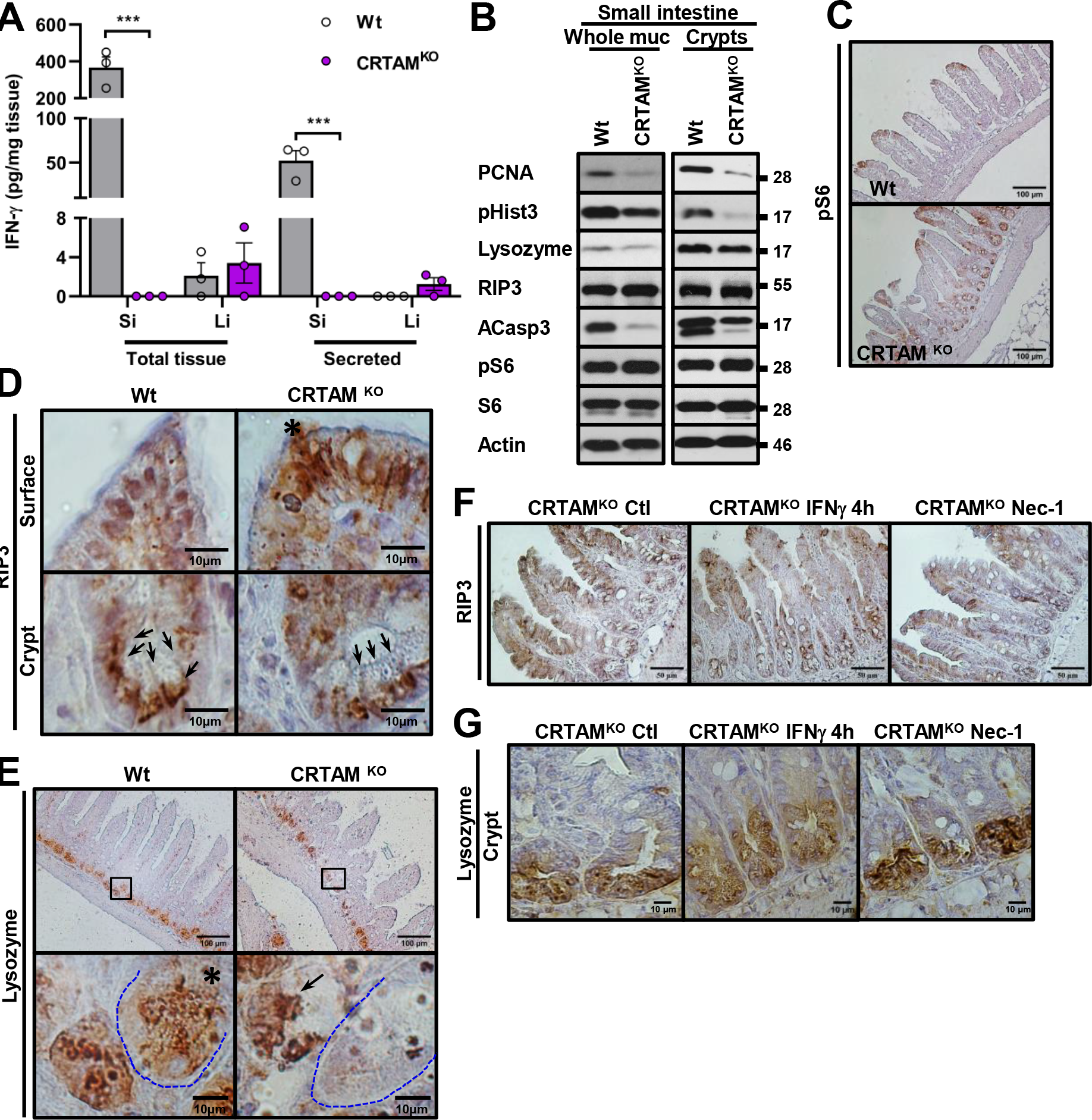
Mucosal IFN-γ is constitutively released in Si but not in Li. A) Quantification of IFN-γ was carried out in small (Si) and large intestine (Li) of Wt and CRTAM^KO^ mice. Whole mucosa or conditional media collected from unstimulated mucosa was tested for IFN-γ using a Biolegend LEGENDplex assay. 3 different animals were evaluated. B) PCNA, pHist3, Lysozyme-1, RIP3, ACasp3, pS6 and S6 were evaluated by western blotting whole mucosa and isolated crypts of small intestine from Wt and CRTAM^KO^ mice. Actin was used as loading control. n=4 independent experiments. C) pS6 was evaluated by IHC in 4 μm paraffin embedded sections of small intestine of Wt and CRTAM^KO^ mice. n= 6 animals per group. Scale bars: 100*μ*m. n= 6 animals per group. D) RIP3 was evaluated by IHC in 4 μm paraffin embedded sections of small intestine of Wt and CRTAM^KO^ mice. Upper panel= Amplification of the villus surface. Lower panel= Amplification of the crypts of Lieberkühn. Arrowheads= necrosomes RIP3+. Scale bars:10*μ*m. n= 6 animals per group. E) Lysozyme-1 was evaluated by IHC in 4 μm paraffin embedded sections of small intestine of Wt and CRTAM^KO^ mice. Upper panel= low magnification. Lower panel= high magnification of square box presented in upper panel. Asterisk=normal looking PC. Arrow=PC with granularly condensed lysosyme-1. C= Crypts of Lieberkühn with atrophic PC lysozyme-1-. Discontinuous line surrounds IEC located at the crypt base. Scale bars: 100*μ*m and10*μ*m. n= 6 animals per group. F) RIP3 was evaluated by IHC in 4 μm paraffin embedded sections of small intestine of Wt and CRTAM^KO^ mice in control conditions or injected with IFN-γ or necrostatin-1. Mice were euthanized 4 post-cytokine and necrostatin-1 injection. n= 6 animals per group. Scale bars: 50*μ*m. n= 6 animals per group. G) Lysozyme-1 was evaluated by IHC in 4 μm paraffin embedded sections of small intestine of CRTAM^KO^ mice in control conditions or injected with IFN-γ or necrostatin-1. Mice were euthanized 4 post-cytokine and necrostatin-1 injection. n= 6 animals per group. Scale bars: 10*μ*m. n= 6 animals per group. Data are shown as mean ± SEM and are pooled from 3 independent experiments. P values were calculated using one-way analysis of variance with the Tukey post hoc test (A). ***p < 0.001.

pS6 or RIP3 levels were similar in whole mucosa and IEC of Si amongst CRTAM^KO^ and Wt mice (**Figure 6B)**. However, active mTORC1 was mainly detected in surface IEC of Wt mice while in CRTAM^KO^ animals the active kinase was also observed at the crypt bottom (**Figure 6C)**. Assembled necrosomes RIP3^+^ were detected in IEC along the crypt- villi axis in both animals though; crypt base IEC necrosomes+ diminished in CRTAM^KO^ mice (**Figure 6D**). In contrast, surface IEC displaying necrosomes RIP3^+^ notoriously increased when CRTAM is lacking (**Figure 6D**). Furthermore, lysozyme-1 protein levels and lysozyme^+^ cells decreased in the Si of CRTAM^KO^ mice; nevertheless, the process was not a generalized event, but rather a regionalized occurrence (**Figure 6B, E**). However, lysozyme-1 was more concentrated in IEC of CRTAM^KO^ mice when compared with IEC of Wt animals (**Figure 6E**). Furthermore, fecal lysozyme-1, an indicative of PC secretion towards the intestinal lumen^32^ was reduced in CRTAM^KO^ mice (**Supplementary figure 6A**). Concerning intestinal ECP (pHist3, PCNA) and apoptosis (ACasp3), both processes were reduced in the Si of CRTAM^KO^ mice (**Figure 6B**). Consequently, the villi of CRTAM^KO^ mice displayed a length shortening (**Supplementary figure 6B**). Considering these results, we next evaluated the effect of IFN-γ and necrostatin-1 in the regulation of PC of CRTAM^KO^ mice. As expected, IFN-γ treatment stimulated the condensation of RIP3 in crypt IEC and augmented the secretion of lysozyme-1 into the crypt-lumen while depleting the cytosolic pool of the protein (**Figure 6F-G**). In contrast, necrostatin-1 administration depleted RIP3 in crypt IEC and induced lysozyme-1 accumulation in cytosolic granules (**Figure 6F-G**). No changes in PC cytoarchitecture were induced by any of the treatments (**Supplementary figure 6C**).

In the Li, the changes in IEC homeostasis were consistent with the spontaneous secretion of IFN-γ in this organ. For instance, active mTORC1 increased in colonocytes of CRTAM^KO^ mice along the crypt-axis (**Supplementary figure 6D**), ECP was reduced, and apoptosis augmented (**Supplementary figure 6E**), although, the colonic crypt height remains unaltered (**Supplementary figure 6G**). Additionally, Reg4 decreased in colonocytes of CRTAM^KO^ mice since its luminal secretion was enhanced (**Supplementary figure 6A, E**). Surprisingly, IFN-γ apparently induced lysozyme-1^+^ expression in colonic IEC of CRTAM^KO^ mice (**Supplementary figure 6F**) and Wt animals (**Supplementary figure 5C**). These results confirm that mucosal IFN-γ modulates IEC homeostasis by controlling PC/PLC function in the gut.

### PC metaplasia during colitis

mTORC1 activation and massive death of colonocytes after DSS treatment (**Figure 7A, Supplementary 7A-E**) are attributable to the colitogenic milieu generated by mucosal cytokines, including IFN-γ^27, 40^. Thus, we evaluated PC/PLC functions in the Li of colitic mice. In control conditions, lysozyme-1 was detected in Si and Reg4 in Li, mainly in the distal region. However, after 6d of treatment lysozyme-1 was found in the distal region of the Li of ∼50% of the samples, indicating PC metaplasia **(Figure 7B)**. IHC analysis corroborated the presence of IEC lysozyme-1^+^ in the colonic mucosa of DSS-treated mice (**Figure 7C)**. Likewise, Reg4 augmented in the distal region of DSS-treated mice. However, in proximal colon increased at 3d post-treatment and was reduced by day 6. Additionally, the fecal content of 1ysozyme-1 and Reg4 augmented in colitic mice, indicating degranulation of PC/PLC. Necrostatin-1 administration reduced the fecal content of both molecules (**Figure 7D-E**). According to these findings, Reg4 diminished in colitic colonocytes but was enriched following necrostatin-1 treatment (**Figure 7F**). These results suggest hypersecretion of PC/PLC in a colitogenic environment. Therefore, we analyzed the proliferation marker PCNA in the Li of colitic mice that were injected with carrier alone or necrostatin-1. DSS-induced colitis resulted in PCNA reduction and the administration of necrostatin-1 further enhanced the process (**Figure 7G)**, indicating that PC/PLC degranulation stimulated IEC proliferation in the colitic mucosa. In conclusion, necroptosis increases PC/PLC secretion to favorably control IEC proliferation in the Li of mice treated with DSS.

**Figure 7.**
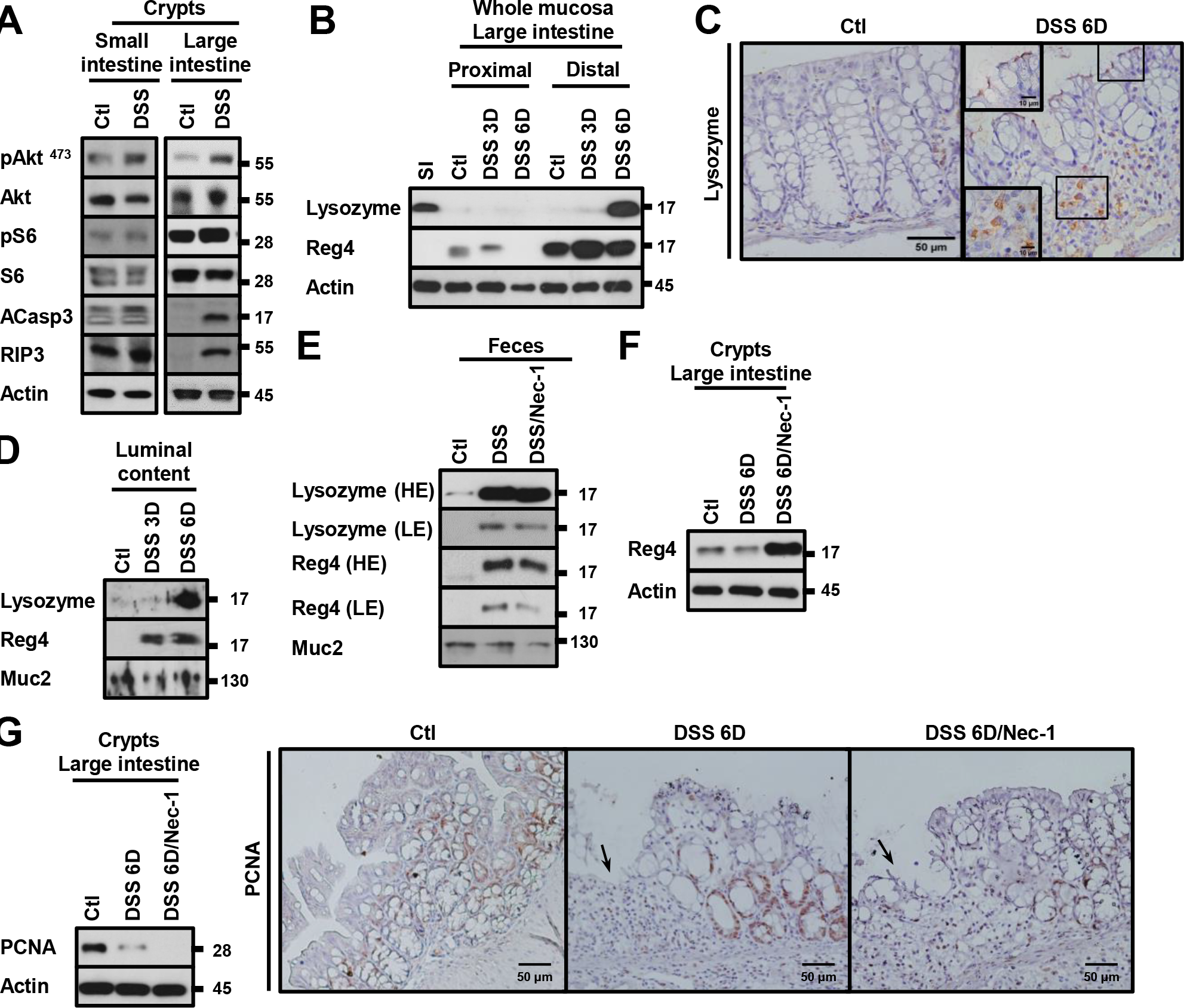
IFN-γ induces PC metaplasia and cell proliferation in distal colon. A) pAkt^473^, Akt, pS6, S6, ACasp3 and RIP3 were evaluated by western blotting isolated crypts of small and large intestine obtained from C57BL/6J mice received water alone or 2.5% DSS dissolved in drinking water for 6d. Actin was used as loading control. Representative blot is shown. n=6 independent experiments. B) Lysozyme-1 and Reg4 were analyzed by Western blotting whole mucosa of large intestine obtained from control and DSS treated animals. Small intestine samples were used as positive control for Lysozyme-1. 2.5% DSS treatment was carried out for 3 and 6d. Actin was used as loading control. Representative blot is shown. n=6 independent experiments. C) Lysozyme-1 was evaluated by IHC in 4 μm paraffin embedded sections of large intestine from DSS-treated mice. Mice were euthanized after 6d of treatment. Inset= Amplification of colonocytes Lysozyme-1+. n=3 animals per group. Scale bars:50*μ*m, 10*μ*m. D) Luminal secretion of Lysozyme and Reg4 was analyzed in feces from control and DSS treated mice. 2.5% DSS treatment was carried out for 3 and 6d. A non-specific band detected by Mucin-2 antibody was used as loading control. Representative blot is shown. n=3 independent experiments. E) Luminal secretion of Lysozyme and Reg4 was analyzed in feces from control, DSS and DSS/Nec-1 (4.5 mg/Kg, i.p.) treated mice. 2.5% DSS treatment was carried out for 6d. Necrostatin-1 was administered 2 h before animals were euthanized. A non-specific band detected by Mucin-2 antibody was used as loading control. Representative blot is shown. n=3 independent experiments. F) Reg4 protein was analyzed by western blotting isolated crypts large intestine from control, DSS and DSS plus Nec-1 treated mice. Necrostatin-1 was administered 2 h before animals were euthanized. Actin was used as loading control. Representative blot is shown. n=3 independent experiments. G) PCNA was evaluated by western blotting and IHC in the large intestine of control, DSS and DSS/Nec-1 treated mice. Actin was used as loading control. Representative blot of n=3 independent experiments is shown. Hyperproliferative areas bordering ulcerated regions (arrows) of DSS and DSS/Nec-1 are shown. Bar= 50 mm.

**Figure 8.**
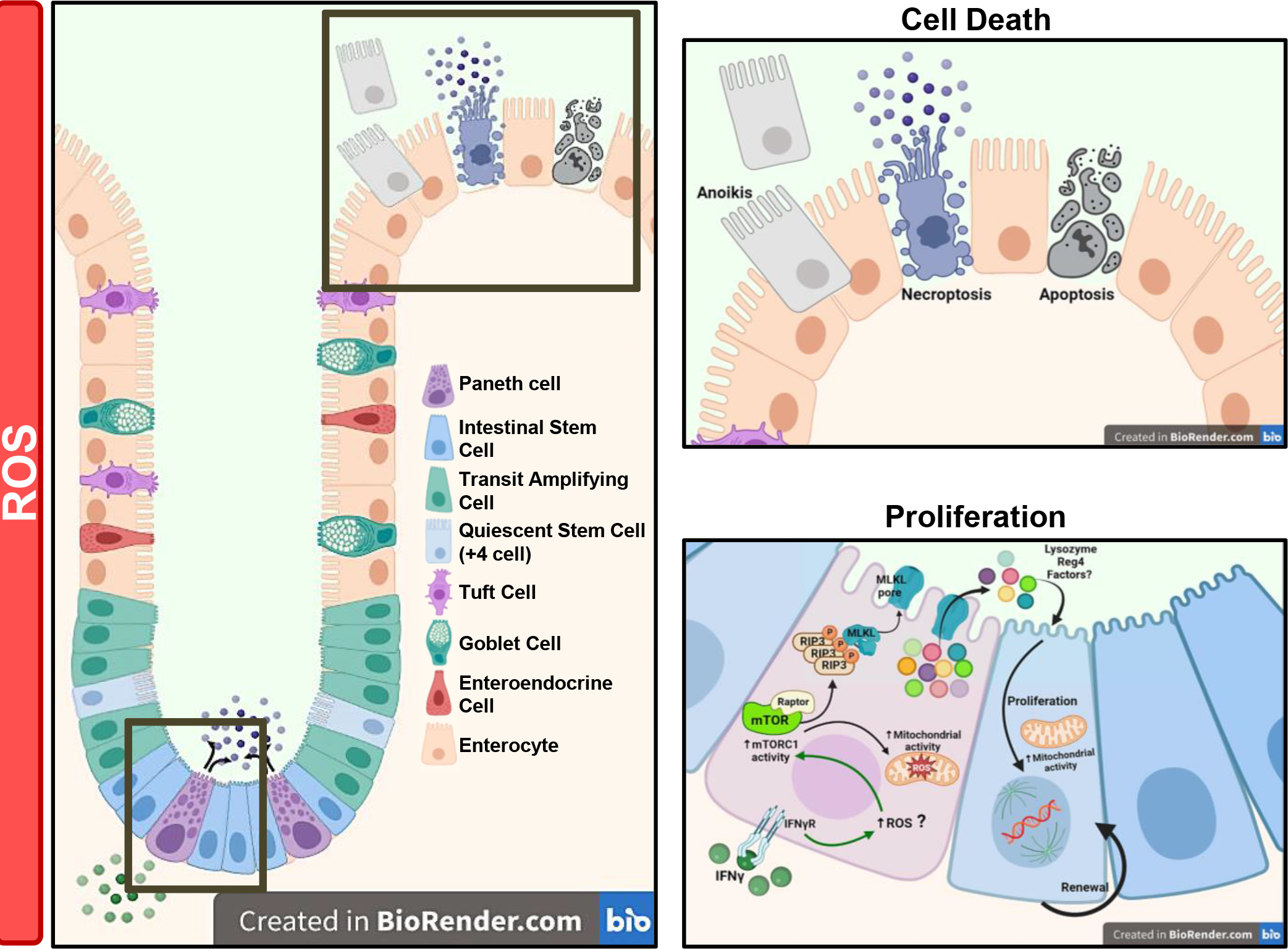
Graphical Abstract. In PC at the crypt of Lieberkühn, IFN-γ induces mitochondrial activation which causes ROS overproduction and mTORC1 activation. Active mTORC1 produces necroptosis to allow degranulation of PC. The release of the PC intracellular contents stimulates proliferation in neighbored cells. Similar events are triggered in PLC at the bottom of the colonic crypts.

## DISCUSSION

Overall, our studies establish a role for the necroptosis machinery in maintaining IEC homeostasis through controlling PC/PLC degranulation. Additionally, our findings demonstrate the participation of colonic IFN-γ in the development of PC metaplasia. PC genesis downstream of IFN-γ could be pathophysiologically relevant if assuming that PC positively regulate IEC proliferation that could explain why PC metaplasia associates with colon cancer and polyposis development in IBD patients^41–43^. Still, given its multifaceted functions in gut physiology we cannot ignore that colonic PC/PLC can mitigate a broader problem. For example, increased PC/PLC degranulation could regulate IEC differentiation/survival^44, 45^, the response to mucosal immunomodulatory signals^46, 47^ and/or induce a defense mechanism that protects the injured mucosa from harmful pathogens^48^. Although conceivable, some of those scenarios can also be disadvantageous for example, over-discharging antimicrobial products by PC/PLC will disturb bacterial load and/or its diversity causing dysbiosis^49–51^. Thus, PC/PLC dysfunction can influence IBD pathogenesis at different levels.

PC genesis, death and function has been extensively studied but remain controversial. However, we compellingly demonstrate the implication of IFN-γ in the process. For instance, IFN-γ expands IEC precursors and activates mTORC1 in the stem cell niche two processes that mediates PC genesis^14, 52, 53^. Further, the cytokine promotes lysozyme-1 expression in colonocytes during PC metaplasia. Thus, undoubtedly IFN-γ influences PC genesis. Additionally, it has been established that IFN-γ stimulates PC death^13^. However, that could be only one side of the story. In fact, in PC undergoing necroptosis the membrane pores will consent the leakage of hydrophilic molecules^54, 55^ that stimulate proliferation, differentiation and cell lineage commitment in neighboring cells^45, 56^. A cell mechanism of such importance must be perfectly controlled, and numerous reports provide evidence in that regard. For example, PC rapidly degranulate and replenish their granule contents^31, 57^. Additionally, PC quickly regenerate and even proliferate/expand in proinflammatory environments^31, 58^. On the contrary, during chronic inflammation or with over-stimulatory signals PC quickly die^13, 59, 60^. Then, multiple mechanisms are intended to modulate PC function.

Signaling pathways driving ECP following IFN-γ administration support the singularity of the system. For example, it was surprising but not extraordinary that PC discharging created a *β*-catenin independent pro-proliferative environment a scenario that could be circumscribed to interferons in general^61–63^. Then, any event inducing interferons secretion will tentatively exploit this mechanism to change IEC homeostasis. Whether this process creates an adaptive or deleterious mechanism must be carefully investigated.

Degranulation of PC/PLC IFN-γ-dependent identifies an unclear but direct connection between mitochondrial stress, ROS, mTORC1 and necroptosis and considering its biological relevance the topic is worth studying. Even though we cannot fully clarify the relationship between the processes, we can certainly assure that the execution of necroptosis follows a similar design to the one described for pyroptosis, and involves mitochondrial hyperactivation/stress, ROS overproduction, and a systematic inhibition of autophagy^7, 64^. Then, ROS/mTORC1 signaling could stimulate necroptosis by inhibiting autophagy and allowing the stabilization of the proteins forming the membrane pores^65–67^. Additionally, mTORC1 phosphorylates and inactivates the Ser/Thr kinase unc-51 like kinase 1 (ULK1) a known inhibitor for necroptosis and pyroptosis^68, 69^. Thus, mTORC1 could be involved in multiple steps during the process. Hence, considering that pyroptosis and necroptosis are highly similar, it is understandable why both processes share a function: the release of intracellular contents targeting neighboring cells.

## Acknowledgments.

The research was partially supported by the sectorial funding for research and education via the grant for basic science from the National Council of Science and Technology (Conacyt) A1-S-20887 (P.N.). The Confocal Microscopy Unit provided confocal microscopy facilities at the Cell Biology Department of CINVESTAV-IPN (Conacyt grant: 300062 to F.G.S.). We want to thank Norma Trejo, Dolores Martín-Tapia, and Juan Unzueta for their technical assistance. M.d.R.E.-G., J.R.D.l.T-B., M.A.H-C. contributed equally to this work. Both J.R.D.l.T-B. (CVU 954035) and M.d.R.E.-G. (CVU 860508) have received CONACYT scholarships. The authors declare that there are no known conflicts of interest associated with this publication and there has been no significant financial support for this work that could have influenced its outcome.

## Notes

### Competing Interest Statement

The authors have declared no competing interest.

