## Supplementary Figures for "Paneth and Paneth-like cells undergoing necroptosis fuel intestinal epithelial cell proliferation following IFN-γ stimulation"

### Supplementary figure 1.

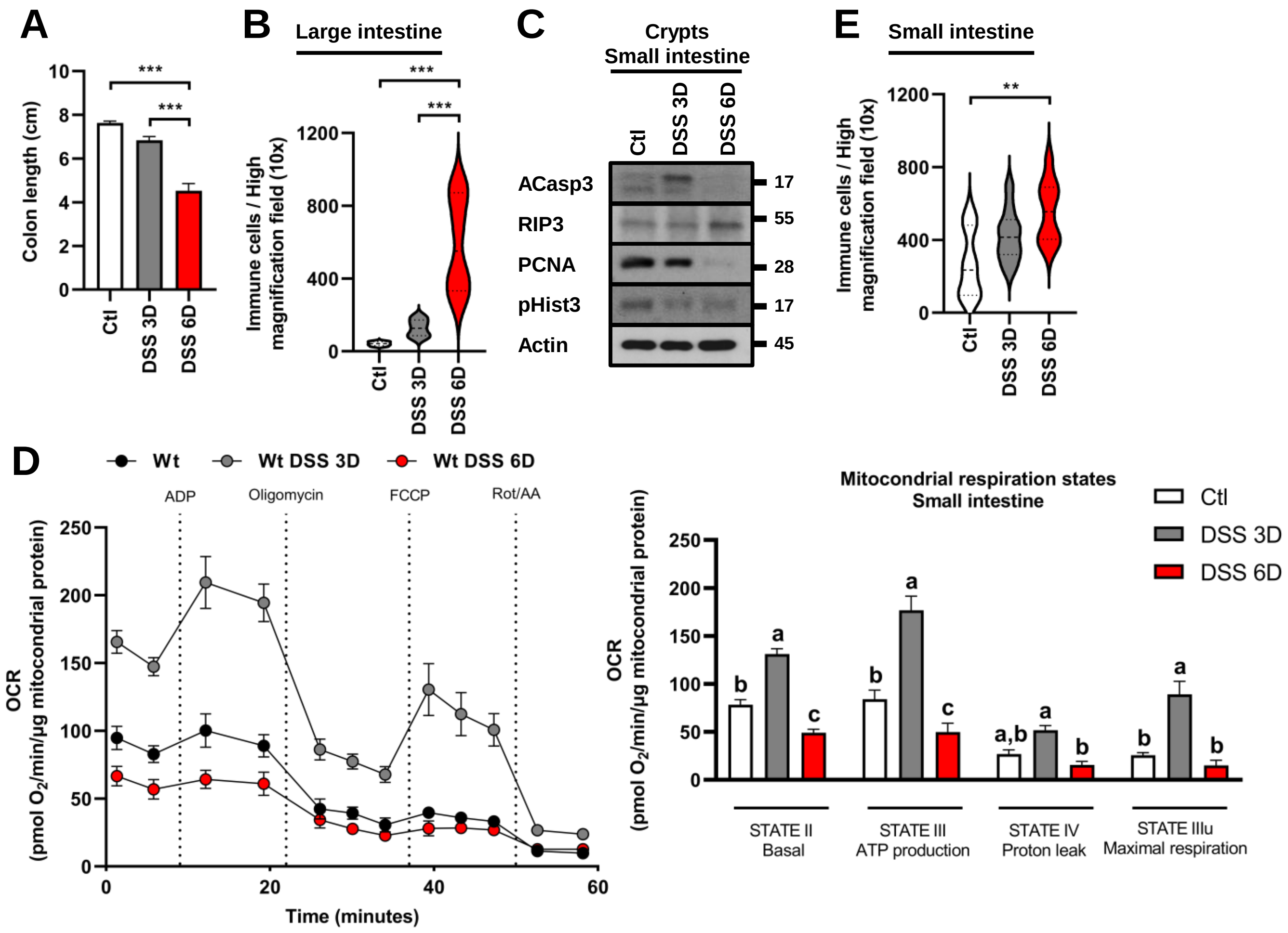

**Supplementary figure 1. Apoptosis and necroptosis in IEC cells.**

A) Graph displaying colon length from control and DSS treated animals. 2.5% DSS treatment was carried out for 3 and 6 days. n=6.

B) Immune cell infiltration was evaluated in the colonic mucosa of control and DSS treated animals. 2.5% DSS treatment was carried out for 3 and 6d. 6 different animals per condition were evaluated.

C) ACasp3, RIP3, PCNA and pHist3 were evaluated by western blotting isolated crypts of small intestine harvested from control and DSS treated animals. 2.5% DSS treatment was carried out for 3 and 6d. Actin was used as loading control. Representative blot of n=6 independent experiments is shown.

D) Oxygen consumption rate (OCR) in isolated mitochondria from small intestine of control and DSS (2.5%)- treated mice after 3 and 6d of treatment. OCR was measured at basal conditions and following ADP, oligomycin, FCCP and antimycin A/rotenone injection. Respiration states were calculated by subtracting OCR values after antimycin A/rotenone administration. n=4-8 mice per group.

E) Immune cell infiltration was evaluated in the small intestine mucosa of control and DSS treated animals. 2.5% DSS treatment was carried out for 3 and 6d. 6 different animals per condition were evaluated.

Data are shown as mean ± SEM and are pooled from 3 independent experiments. P values were calculated using one-way analysis of variance with the Tukey post hoc test (A, B, D, E). $^{**}p < 0.01$ $^{***}p < 0.001$. Mean values with different lowercase letters show statistical differences between each other were a>b>c (D).

### Supplementary figure 2.

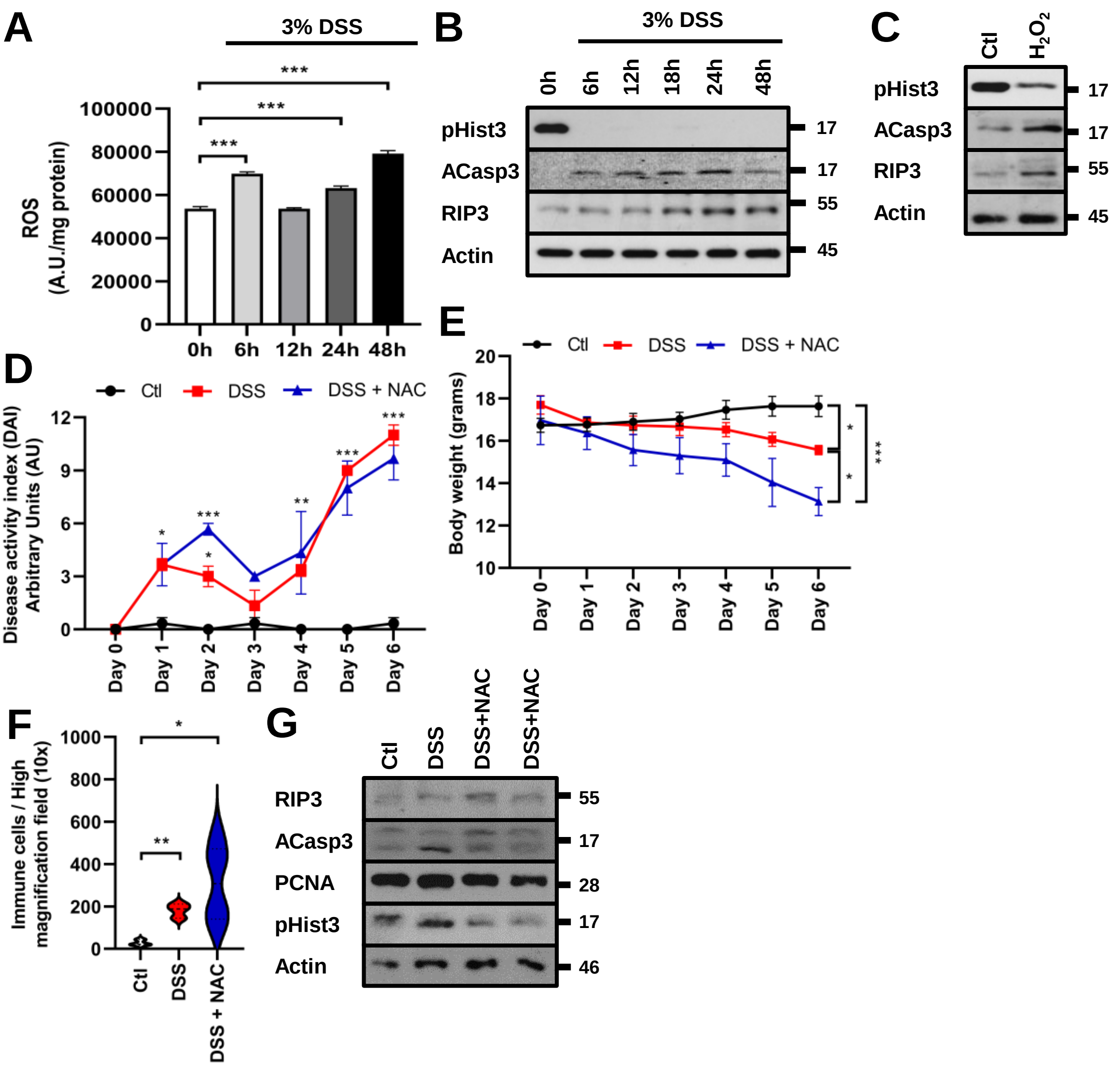

**Supplementary figure 2. NAC treatment attenuates DSS-induced damage.**

A) Quantification of ROS in SW480 cells treated with 3% of DSS. ROS were measured in cell lysates using the fluorescent CellROX™ assay. DSS treatment was carried out for 0h, 6h, 12h, 24h and 48h. A histogram displaying the mean of 3 independent experiments is shown.

B) pHist3, ACasp3 and RIP3 were assessed in cell lysates obtained from SW480 cells. 3% DSS treatment was carried out for 0h, 6h, 12h, 24h and 48h. Actin was used as loading control. n=3 independent experiments.

C) Representative Western blot analysis shows pHist3, ACasp3 and RIP3 in cell lysates obtained from SW480 cells. 20µM $H_2O_2$ treatment was carried out for 4hours. Actin was used as loading control. n=3.

D) Disease activity index (DAI) of C57BL/6J mice treated with water (Ctl), 2.5% DSS, or 2.5% DSS plus 100 mg/Kg N-Acetyl-L-cysteine (DSS/NAC) for 6d. NAC was administered daily via i.p. Disease activity index is expressed in arbitrary units. n=6 animals per group.

E) Graph displaying body weight loss of 2.5% DSS-treated mice. Animals were administered orally with DSS dissolved in drinking water or DSS plus 100 mg/Kg N-Acetyl-L-cysteine (DSS/NAC). Body weight was assessed daily prior to NAC administration (i.p.). n=6 animals per group.

F) Immune cell infiltration was evaluated in the colonic mucosa of control, DSS and DSS/NAC treated animals. n=6 animals per group. 2.5% DSS treatment was carried out for 6d. NAC administration was carried out i.p. 6 different animals per condition were evaluated.

G) RIP3, ACasp3, PCNA and pHist3 were evaluated by western blotting isolated crypts of large intestine harvested from control, DSS and DSS/NAC treated animals. Wt mice were administered orally with 2.5% DSS for 3d. NAC (100 mg/Kg) was administered daily via i.p. Actin was used as loading control. n=4 independent experiments.

Data are shown as mean ± SEM and are pooled from 3 independent experiments. P values were calculated using two-way ANOVA with Sidak's multiple comparison test (D, E) and one-way analysis of variance with the Tukey post hoc test (A, F). *$p < 0.05$; **$p < 0.01$; ***$p < 0.001$.

### Supplementary Figure 3.

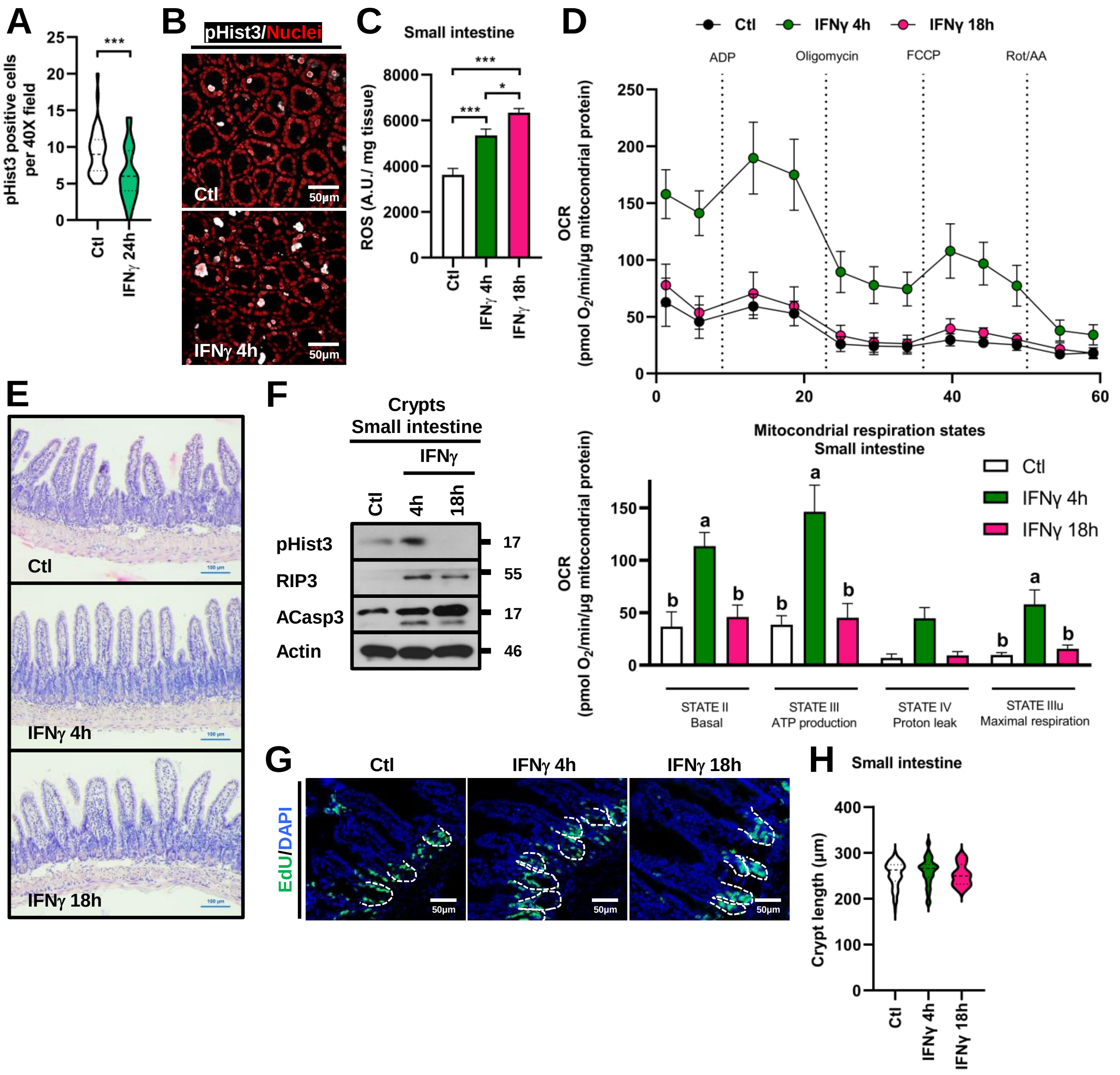


**Supplementary Figure 3. IFNγ regulates proliferation in intestinal epithelial cells.**

A) Quantification of pHist3$^{+}$ cells per high magnification field (40X) in the large intestine of C57BL/6J mice injected i.p. with MSA or IFN-γ (2.5 mg/kg). Mice were euthanized 24h post-cytokine injection. n=6 independent experiments.

B) Immunofluorescence staining for pHist3 (white) in colonic cryosections of control and IFN-γ treated mice. Nuclei=red. Bar=50µm. n=6. A representative image is displayed.

C) Quantification of ROS in small intestine harvested from C57BL/6J mice injected i.p. with MSA or IFN-γ (2.5 mg/kg). ROS were measured in whole mucosal cell lysates using the fluorescent CellROX™ assay. Cytokine treatment was carried out for 4 or 18h. A histogram displaying the average n of 3 independent experiments is shown.

D) Oxygen consumption rate (OCR) in isolated mitochondria from small intestine of C57BL/6J mice injected i.p. with vehicle (Mouse Serum Albumin, MSA) or IFNγ (2.5 mg/kg) for 4 or 18h post-injection. OCR was measured at basal conditions and following ADP, oligomycin, FCCP and antimycin A/rotenone injection. Respirations states were calculated by subtracting OCR values after antimycin A/rotenone administration. n=4-8 mice per group.

E) H&E staining obtained from small intestine of control and IFN-γ treated mice. Mice were euthanized 4 or 18h post interferon-injection. Bar= 100µm. Representative images are shown. n=6 animals per group.

F) pHist3, RIP3 and ACasp3 were assessed by western blotting isolated crypts obtained from small intestine of C57BL/6J mice injected i.p. with MSA or IFN-γ (2.5 mg/kg). Mice were euthanized 4 or 18h post-cytokine injection. Actin was used as loading control. n=6 independent experiments.

G) Incorporation of EdU (green) was analyzed in cryosections of small intestine by confocal microscopy. C57BL/6J mice were injected i.p. with MSA or IFN-γ (4 and 18h, 2.5 mg/kg). Representative images are shown. Nuclei=blue. A discontinuous white line marks crypt bottom. Bar=50µm.

H) Graph showing the crypt length of the small intestine tissue harvested from each group. C57BL/6J mice were injected i.p. with MSA or IFN-γ (4 and 18h, 2.5 mg/kg). n=6 animals per group.

Data are shown as mean ± SEM and are pooled from 3 independent experiments. P values were calculated using *t*-test (A), or one-way analysis of variance with the Tukey post hoc test (C, D, H). $^{*}p < 0.05$; $^{**}p < 0.01$; $^{***}p < 0.001$. Mean values with different lowercase letters show statistical differences between each other were a>b>c (D).

### Supplementary Figure 4.

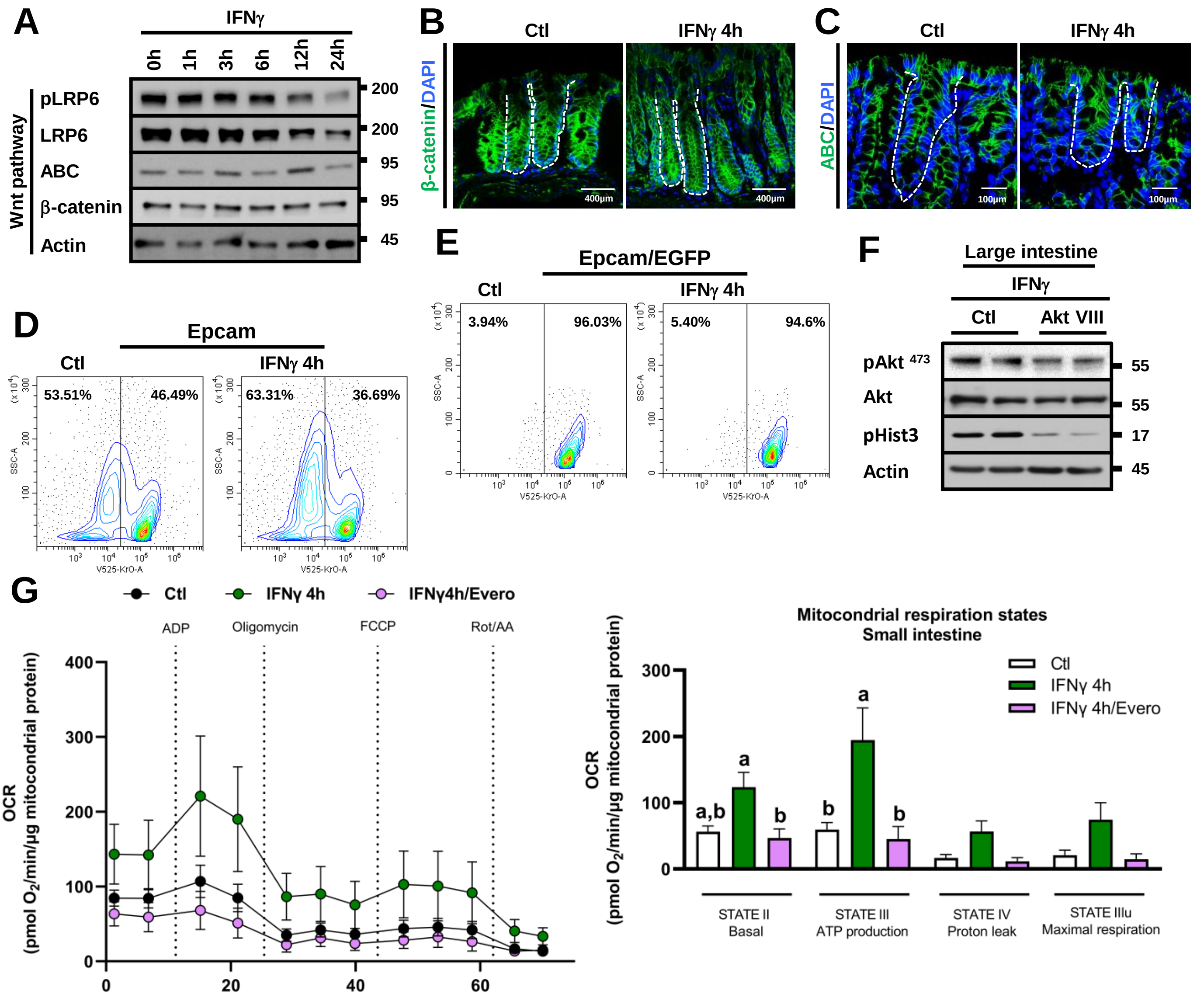

**Supplementary Figure 4. IFNγ increases cell proliferation in IEC through β-catenin.**

A) pLRP6, LRP6, ABC and β-catenin were assessed by western blotting from whole colonic mucosa obtained from control and IFNγ treated animals. Lgr5-EGFP-IRES-creERT2 "knock-in" reporter mice were euthanized 1, 3, 6, 12 and 24h after cytokine administration. Actin was used as a loading control. Representative blot of n=4 independent experiments is shown.

Immunofluorescence (IF) staining for β-catenin (green) (B) or Active β-catenin (ABC, green) (C) in colonic cryosections of control and IFNγ treated mice. Mice were euthanized 4h post interferon-injection. Nuclei=blue. Discontinuous white line surrounds epithelial cells located at the crypt. Bar=50μm. 6 different animals were evaluated, and a representative image is displayed.

Flow cytometric identification of and EpCAM+ (D) and EGFP+/EpCAM+ (E) cells in preparations pooled from ileum of Lgr5-EGFP-IRES-creERT2 "knock-in" reporter mice. Cells were labeled with EpCAM-PE. Values represent the mean percentage of events falling within their respective quadrants. Small intestine was harvested from mice treated with MSA i.p. (Ctl) or IFNγ (2.5 mg/kg, i.p.) for 4h. n=4.

F) $pAkt^{473}$, Akt and pHist3 were evaluated by western blotting large intestine samples obtained from C57BL/6J mice injected i.p. with IFNγ (2.5 mg/kg) or IFNγ plus Akt VIII (Sigma St Louis, MO, 10 mg/Kg). Mice were euthanized 4h post-injection. Actin was used as a loading control. Representative blot of n=3 independent experiments is shown.

G) Oxygen consumption rate (OCR) in isolated mitochondria from small intestine of C57BL/6J mice injected i.p. with MSA (Ctl), IFNγ (2.5 mg/kg), or IFNγ/Everolimus (1 mg/Kg, i.p.) after 4h of treatment. OCR was measured at basal conditions and following ADP, oligomycin, FCCP and antimycin A/rotenone injection. Respiration states were calculated by subtracting OCR values after antimycin A/rotenone administration. n=4-8 mice per group.

Data are expressed as the mean ± SEM. Data were analyzed by one-way ANOVA followed by Tukey multiple comparison post hoc test. The differences were considered statistically significant at $p < 0.05$. Mean values with different lowercase letters show statistical differences between each other were a>b>c (G).

Supplementary figure 5.

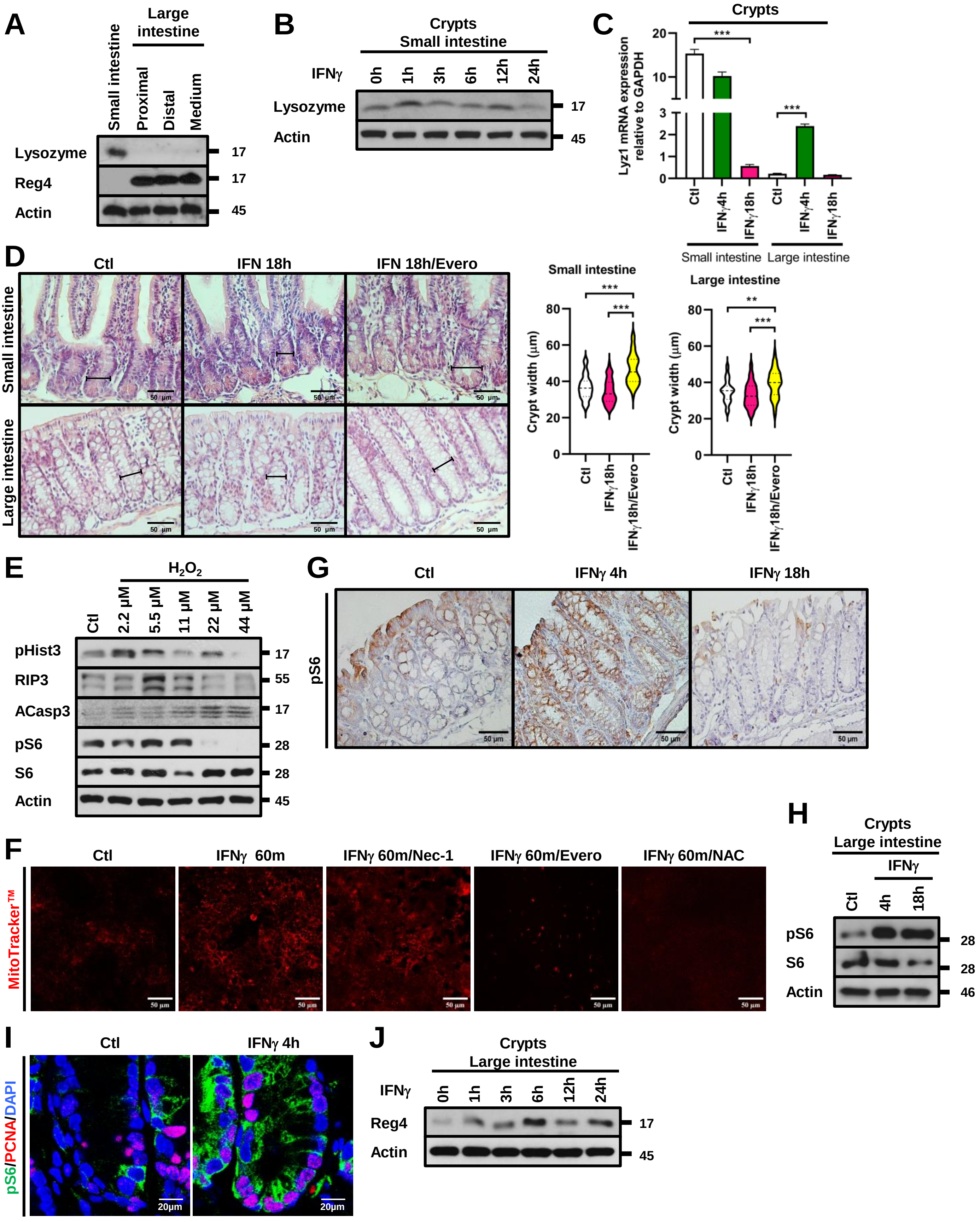

**Supplementary figure 5. mTORC1, Paneth and Paneth-like cell analysis in the gut.**

A) Lysozyme and Reg4 were analyzed by Western blotting whole mucosal samples of small and large intestine obtained from C57BL/6J mice. Actin was used as loading control. n=6 independent experiment.

B) Lysozyme was assessed by western blotting isolated crypts harvested from small intestine of C57BL/6J mice injected i.p. with IFN-γ (2.5 mg/kg). Mice were euthanized 1h, 3h, 6h, 12h or 24h post-cytokine injection. Actin was used as loading control. n=6 independent experiments.

C) mRNA expression of Lyz1 in isolated crypts of small and large intestine from Wt mice treated with MSA i.p. (Ctl) or IFN-γ (2.5 mg/kg, i.p.). Mice were euthanized 4 or 18h post-cytokine injection. Lyz1 expression was normalized to GAPDH.

D) PC morphology and crypt lumen diameter were evaluated in small intestine of control, IFN-γ or IFN-γ/Everolimus treated animals Mice were euthanized 18h post-IFN-γ injection. Everolimus was administered i.p. 30 min before cytokine stimulation. Bar= 50µm. n=6.

E) pHist3, RIP3, ACasp3, pS6 and S6 were evaluated by Western blotting in IEC exposed to different concentrations of $H_2O_2$. SW480 cells were stimulated for 4h. Actin was used as loading control. Representative blot of n=3 independent experiments is shown.

F) Mitochondrial activity was analyzed in SW480 cell monolayers stimulated with IFN-γ using MitoTracker™ Red CMXRos. Confluent monolayers were treated with IFN-γ (100 U/ml), IFN-γ/Necrostatine-1 (80µM), IFN-γ/Everolimus (20µM) or IFN-γ/NAC (100µM). Inhibitors were administered 15 prior cytokine stimulation. IFN-γ-treatment was carried out for 60 minutes. n=3.

pS6 was detected by immunohistochemistry (IHC) (G) and western blotting (H) in IEC at the colonic mucosa of control and IFN-γ (2.5 mg/kg) treated mice. Mice were euthanized 4 or 18h post-cytokine injection. Paraffin embedded sections were 4 µm thick. Isolated crypts were obtained from distal colon. S6 and actin were used as loading controls. Representative blot of n=6 independent experiments is shown.

I) Immunofluorescence (IF) staining for pS6 (green) and PCNA (red) in colonic cryosections of control and IFN-γ treated mice. Mice were euthanized 4h post interferon-injection. Nuclei=blue. Bar=20µm. 6 different animals were evaluated, and a representative image is displayed.

J) Reg4 was assessed by western blotting colonocytes obtained from C57BL/6J mice injected i.p. with IFN-γ (2.5 mg/kg). Mice were euthanized 0h, 1h, 3h, 6h, 12h or 24h post-cytokine injection. Actin was used as loading control. Representative blot of n=6 independent experiments is shown.

Data are shown as mean ± SEM and are pooled from 3 independent experiments. P values were calculated using one-way analysis of variance with the Tukey post hoc test (C,D). **$p < 0.01$; ***$p < 0.001$.

### Supplementary figure 6.

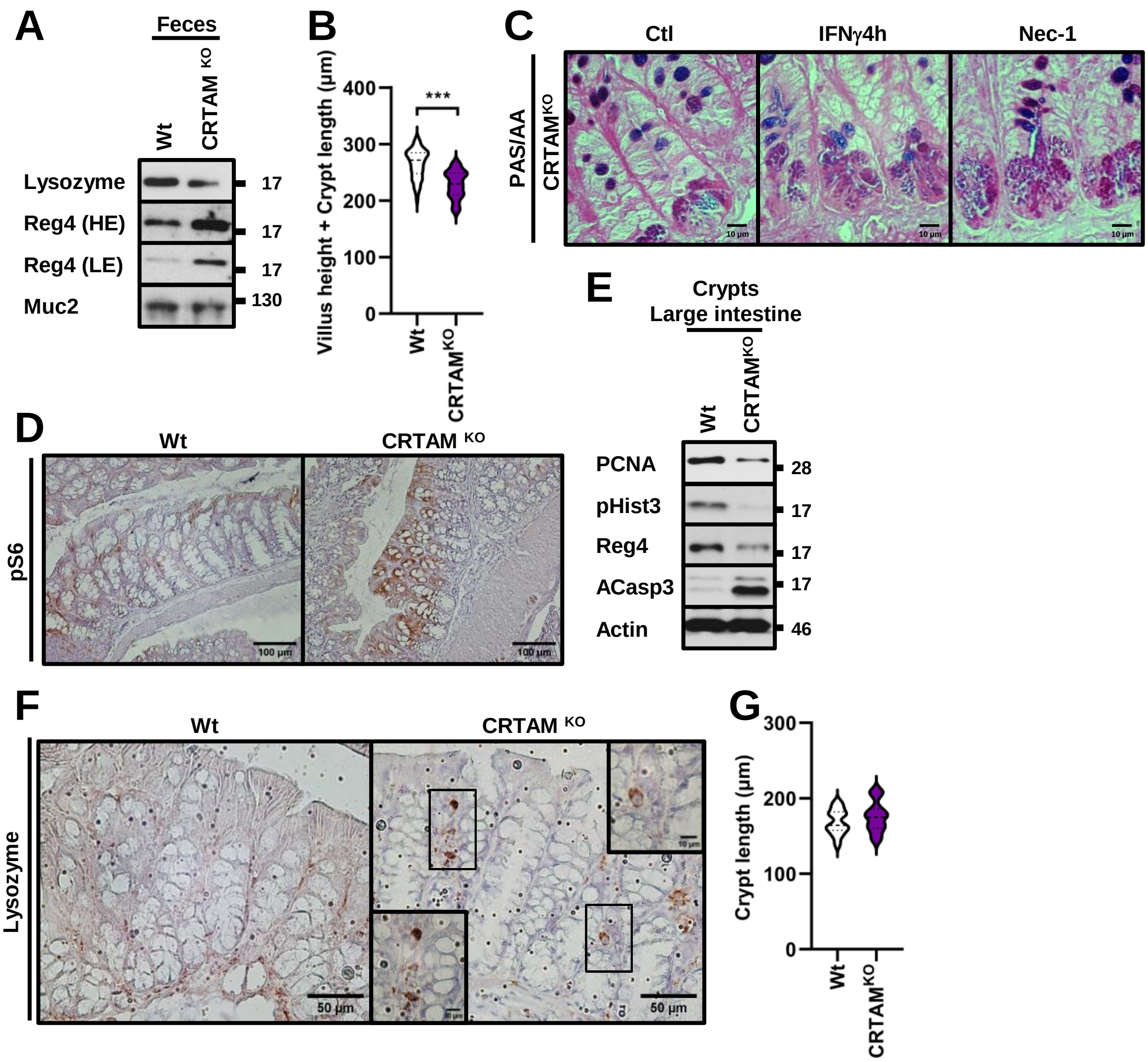

**Supplementary figure 6. Lysosyme-1 is expressed in the colonic mucosa of CRTAM$^{KO}$ mice.**

A) Lysozyme and Reg4 were analyzed in stool samples of unstimulated Wt and CRTAM$^{KO}$ mice. Non-specific band recognized by Muc-2 antibody was used as loading control. Representative blot of n=4 independent experiments is displayed.

B) Histogram showing the villus height plus crypt length of the small intestine units from Wt and CRTAM$^{KO}$ mice. 100 units/structures were analyzed per animal. n=3 per condition.

C) Micrograph of PC of small intestine of CRTAM$^{KO}$ mice in control conditions or injected with IFN-γ or necrostatin-1. PAS/AB staining was carried out in 4 μm paraffin embedded sections of small intestine. Mice were euthanized 4h post-cytokine injection. Bar=10 μm

D) pS6 was assessed by IHC in paraffin embedded sections of large intestine from C57BL/6J (Wt) and CRTAM$^{KO}$ mice. Scale bars: 100μm.

E) PCNA, pHist3, Reg4 and ACasp3 were evaluated by western blotting isolated crypts of large intestine from Wt and CRTAM$^{KO}$ mice. Actin was used as loading control. Representative blot of n=4 independent experiments is shown.

F) Lysozyme was assessed by IHC in paraffin embedded sections of large intestine from C57BL/6J (Wt) and CRTAM$^{KO}$ mice. Inset = Black square internal area. Scale bars: 50μm, 10μm.

G) Graph displaying crypt length of the colonic crypts harvested from each group. 100 units/structures were analyzed per animal. n=3 per condition.

Data are shown as mean ± SEM and are pooled from 3 independent experiments. P values were calculated using *t*-test (B, G) analysis. ***$p < 0.001$.

### Supplementary figure 7.

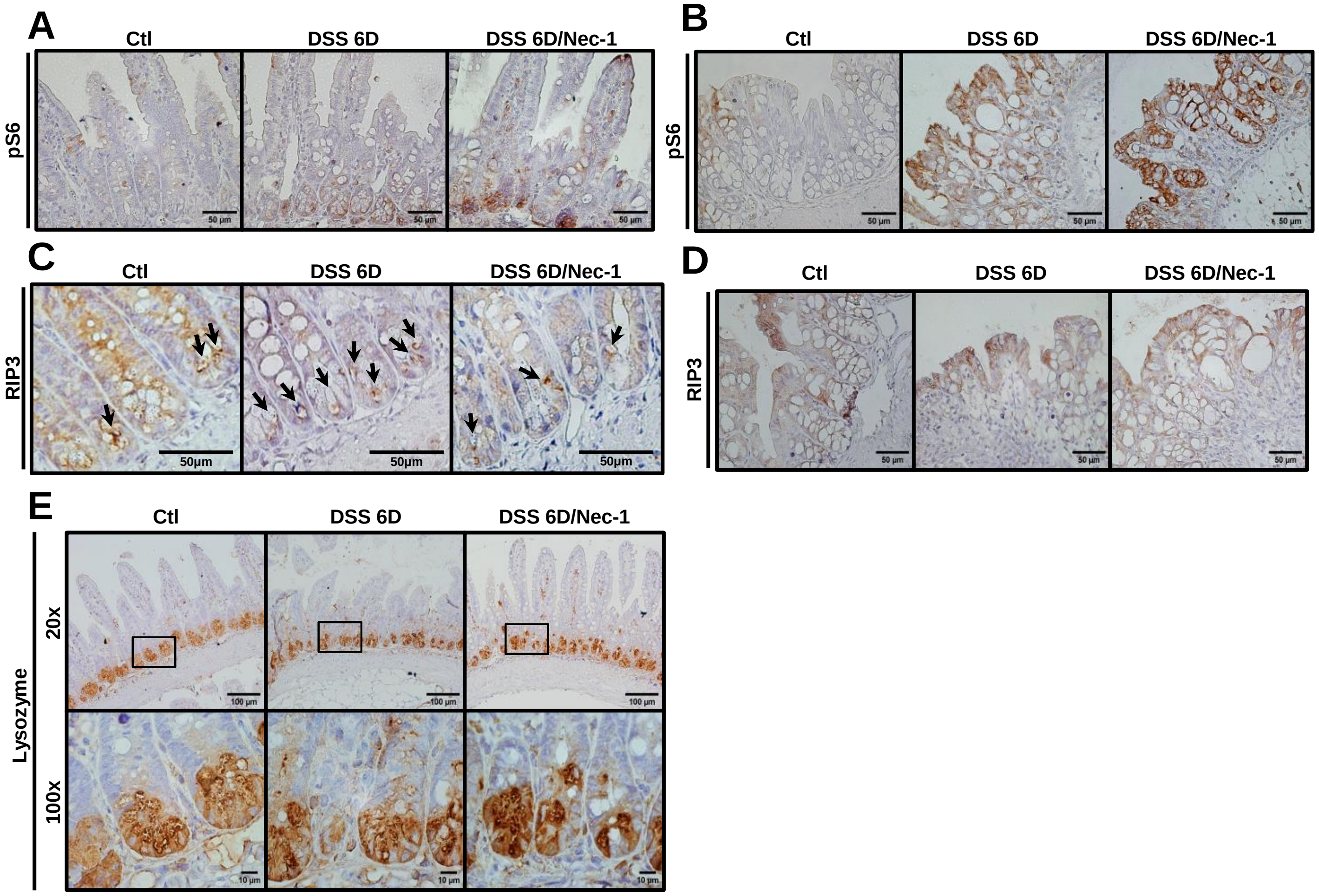


**Supplementary figure 7. Effect of necrostatin-1 in the colitic mucosa.**

pS6, RIP3 and Lyzozime-1 were evaluated by immunohistochemistry (IHC) in paraffin embedded sections of small intestine (A, C and E) and large intestine (B and D) from control, DSS (2.5%) and DSS/ Nec-1 (4.5 mg/Kg, i.p.) treated mice. 2.5% DSS treatment was carried out for 6d. Representative image of n=6 independent experiments is shown. Bar=100µm, 50µm or 10µm.
